# Non-Invasive Perfusion MR Imaging of the Human Brain via Breath-Holding

**DOI:** 10.1101/2023.08.24.554712

**Authors:** J.B. Schulman, S. Kashyap, S.G. Kim, K. Uludağ

**Affiliations:** Department of Medical Biophysics, University of Toronto, Toronto, ON, Canada; Krembil Brain Institute, University Health Network, Toronto, ON, Canada; Center for Neuroscience Imaging Research, Institute for Basic Science, Suwon, Republic of Korea; Department of Biomedical Engineering, Sungkyunkwan University, Suwon, Republic of Korea; Physical Sciences, Sunnybrook Research Institute, Toronto, ON, Canada

**Keywords:** MRI, Perfusion, DSC, Contrast, Deoxyhemoglobin

## Abstract

Dynamic susceptibility contrast (DSC) MRI plays a pivotal role in the accurate diagnosis and prognosis of several neurovascular diseases, but is limited by its reliance on gadolinium, an intravascularly injected chelated metal. Here, we determined the feasibility of measuring perfusion using a DSC analysis of breath-hold-induced gradient-echo-MRI signal changes. We acquired data at both 3T and 7T from ten healthy participants who engaged in eight consecutive breath-holds. By pairing a novel arterial input function strategy with a standard DSC MRI analysis, we measured the cerebral blood volume, flow, and transit delay, and found values to agree with those documented in the literature using gadolinium. We also observed voxel-wise agreement between breath-hold and arterial spin labeling measures of cerebral blood flow. Breath-holding resulted in significantly higher contrast-to-noise (6.2 at 3T vs 8.5 at 7T) and gray matter-to-white matter contrast at higher field strength. Finally, using a simulation framework to assess the effect of dynamic vasodilation on perfusion estimation, we found global perfusion underestimation of 20-40%. For the first time, we have assessed the feasibility of and limitations associated with using breath-holds for perfusion estimation with DSC. We hope that the methods and results presented in this study will help pave the way toward contrast-free perfusion imaging, in both basic and clinical research.

## Introduction

Cerebral perfusion imaging, in addition to its research utility in cognitive neuroscience, allows clinicians to investigate diseases characterized by vascular abnormality, including stroke, cancer, and neurodegenerative disease.^[1],[2],[3],[4]^ Although multiple perfusion imaging techniques exist, including the non-invasive arterial spin labeling (ASL), dynamic susceptibility contrast (DSC) MRI is considered the standard perfusion imaging technique in the clinical domain; here, a bolus of paramagnetic contrast agent is tracked as it passes through the cerebral vasculature, and the associated time course properties reflect underlying tissue perfusion in accordance with the principles of indicator dilution theory.^[5],[6],[7],[8]^ DSC yields perfusion metrics possessing significant research and clinical utility, including the cerebral blood volume (CBV), cerebral blood flow (CBF), and mean transit time (MTT).^[1],[2],[3],[9]^ Of note, all DSC perfusion metrics are technically relative, in that they are dependent on numerous acquisition and analysis parameters, as has been previously documented.^[10],[11]^

While gadolinium (Gd), an exogenous paramagnetic metal, is the standard contrast agent used for DSC, researchers have recently exploited the paramagnetic properties of deoxyhemoglobin (dOHb) as an endogenous contrast alternative^[11],[12],[13],[14],[15],[16]^—in essence, both Gd and dOHb induce T_2_* signal changes that can be exploited in a DSC analysis. However, Gd is limited in that it is invasive, expensive, and toxic in certain patient populations.^[17],[18],[19]^ On the other hand, generating dOHb contrast has thus far relied on gas control systems that modify dOHb concentration in the blood through the induction of hypoxia^[11],[12],[13],[14],[15]^, hyperoxia^[20]^, or hyper/hypocapnia^[16]^. Given the associated cost, set-up time, and expertise required for gas control system utilization, the widespread implementation of dOHb contrast as an alternative to Gd is currently limited.

Unlike hypoxia and hyperoxia, hypercapnia (i.e., an increase in arterial CO_2_) does not directly result in a change to the concentration of dOHb, rather, the CO_2_ bolus induces vasodilation through a process known as cerebrovascular reactivity (CVR), resulting in a subsequent rise of *CBF* and reduction of dOHb in the tissue.^[21]^ Given these inherent mechanistic differences, and the fact that a physiological challenge is being used to assess baseline perfusion, the use of hypercapnia for DSC relies on certain assumptions, including the constancy of oxygen metabolism, and most importantly, the ability to measure an arterial input function (AIF), which represents the input time course of contrast agent, or in this case, vasodilatory agent (see Discussion). Although it may seem counterintuitive to measure baseline perfusion using a physiological challenge (i.e., hypercapnia), it should be noted that previous works within cognitive neuroscience have employed vascular challenges to normalize blood oxygenation level-dependent (BOLD)-based estimates of neuronal activity to venous CBV^[4],[22]^ (see also calibrated BOLD ‘M’ parameter, which scales with baseline *CBV* and is calculated using hypercapnic calibration^[23]^).

From a physiological perspective, the effect of breath-holding is practically equivalent to that of hypercapnia induced by a gas control system.^[21],[24],[25]^ However, there is no study that has investigated whether a DSC analysis pipeline could be used to measure perfusion from signal changes induced by breath-holding. Thus, we developed a breath-hold DSC (bhDSC) approach, where subjects performed eight breath-holds (16 s each separated by 44 s of rest) at 3T and 7T, to investigate whether perfusion metrics could be reliably estimated without the use of Gd or a gas control system.

## Results

Gradient-echo 2D-EPI (GRE-EPI) imaging data were acquired on Siemens 3T Prisma and 7T Terra scanners from ten healthy subjects—each subject was scanned at both field strengths. The subjects were instructed to fixate on a countdown timer projected onto the back of the scanner, which indicated when to breathe regularly and when to perform a breath-hold. The first eight breath-hold boluses were allotted a 72 s window centered at the bolus maximum and subsequently averaged (Figure 1).

**Figure 1.**
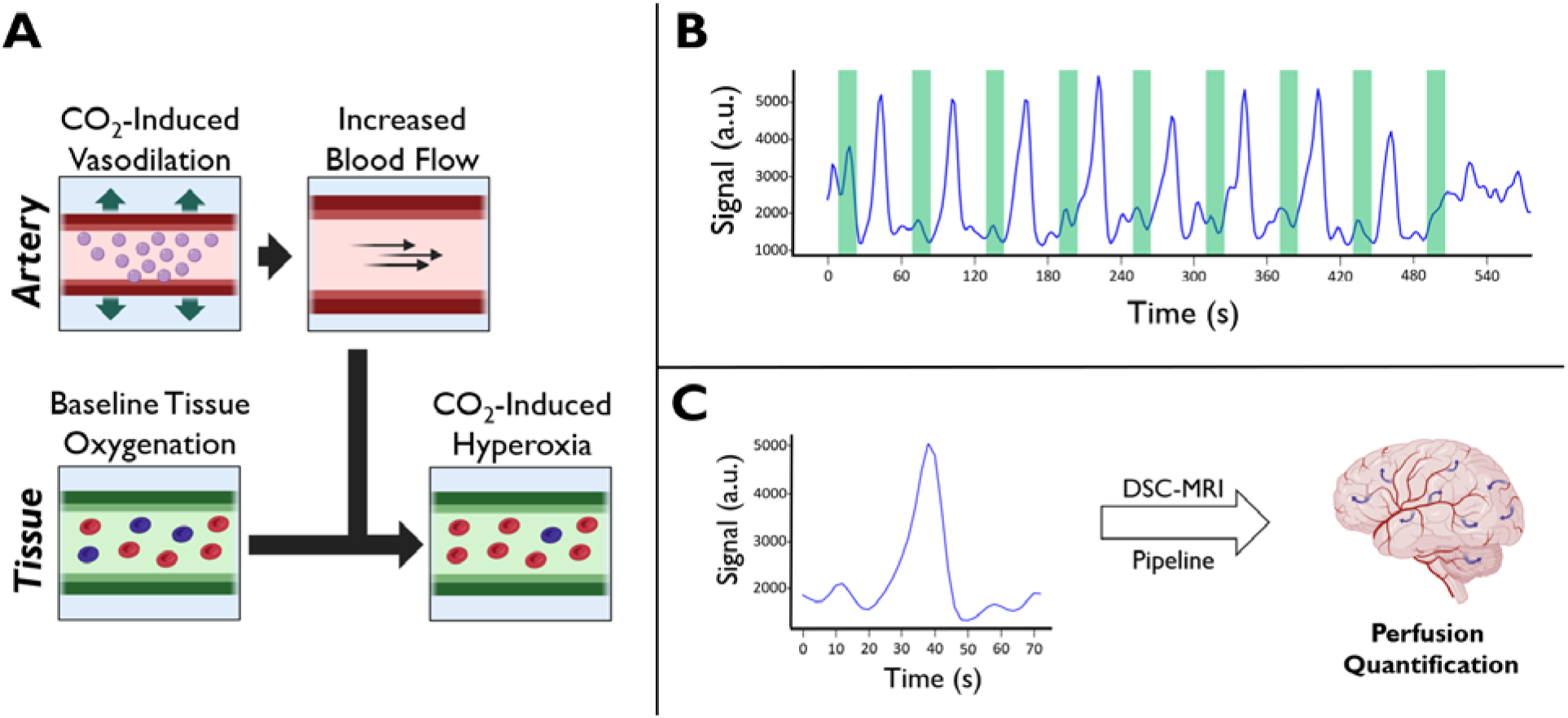
Summary of the Breath-hold Experiment. **A.** Description of the cerebrovascular reactivity phenomenon. At baseline, tissue blood oxygenation is between 60-80%. Increased CO_2_ from breath-holding causes vasodilation in the upstream arterial/arteriolar vasculature, leading to increased blood flow, and subsequently, a decrease in the downstream tissue dOHb concentration (hyperoxia), resulting in a GRE-MRI signal increase. **B.** Signal time course (blue) resulting from nine consecutive breath-holds (green) in a representative venous voxel. **C.** The signal time course resulting from the temporal averaging of the first eight boluses was then used for perfusion estimation.

### Time Course Properties in bhDSC

The temporally averaged signal time courses were then converted to relaxation rate time courses (ΔR_2_*(t)) by normalizing to the pre- and post-breath-hold baselines.

Figure 2 shows the breath-hold-induced ΔR_2_* time courses averaged across all subjects, for arterial (AIF), venous (VOF), gray matter (GM), and white matter (WM) voxels at 3T and 7T. ΔR_2_*(t) bolus dispersion and delay increases from input (AIF), to tissue (GM and WM), and ultimately to output (VOF), as is expected physiologically (Figure 2). Unlike the other time courses, the AIF has a positive ΔR_2_* (to be discussed in Figure 4 and the Discussion section). The bolus magnitude (defined here as the ΔR_2_*(t) integral (AUC)) was found to be ∼1.5 times higher in GM relative to WM. Importantly, breath-holding at 7T resulted in larger bolus magnitudes in the WM and GM (by ∼2 times), artery (by ∼2 times), and vein (by ∼3 times) relative to those at 3T—this is also illustrated in the subject-wise data (Figure S1) and simulation results (Figure S2).

**Figure 2.**
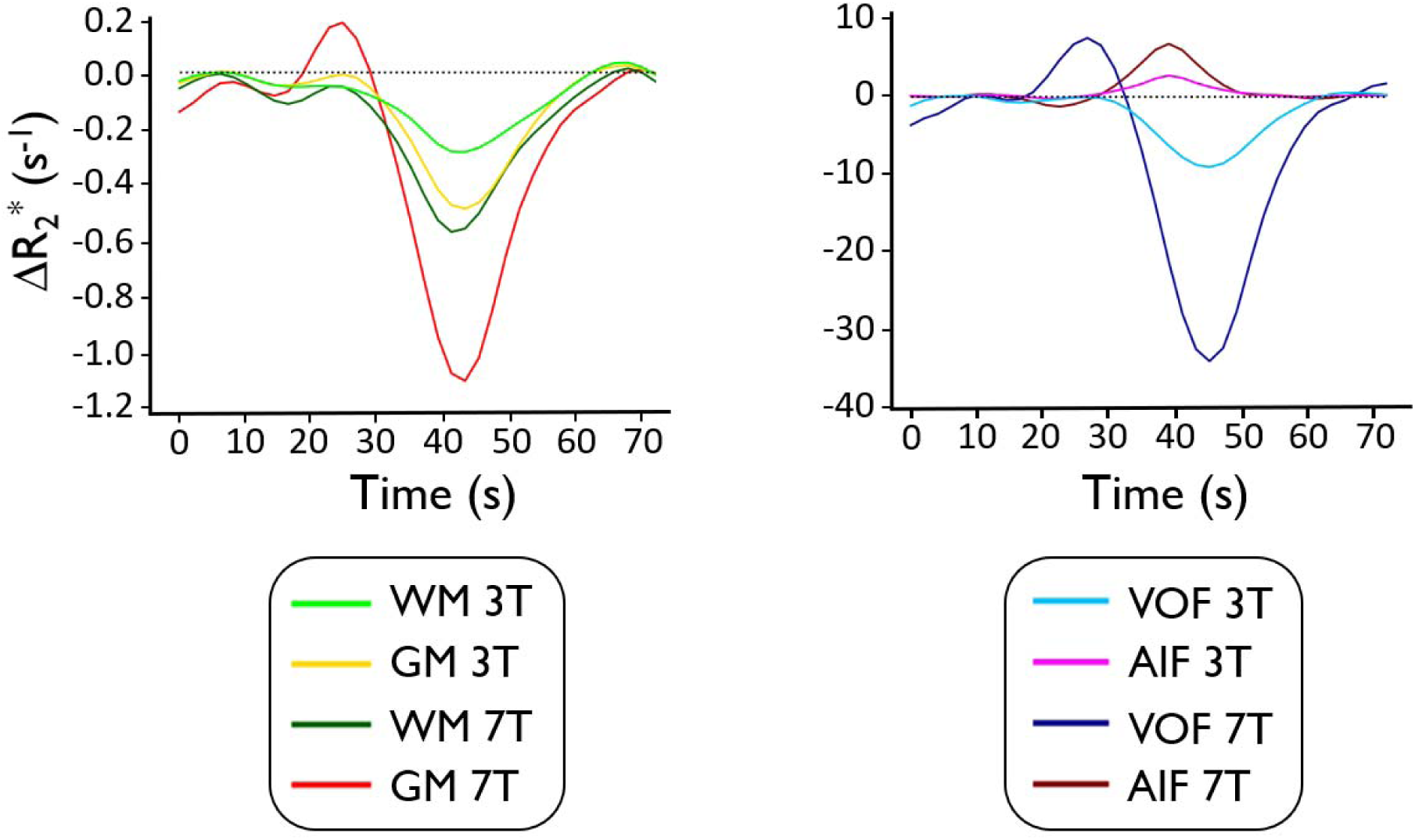
Breath-Hold-Induced Relaxation Rate Time Course Dynamics. Time courses were averaged across subjects for gray matter (GM), white matter (WM), vein (VOF), and artery (AIF) at 3T and 7T.

### Breath-Hold CNR Properties

The contrast-to-noise ratio (CNR) and associated GM-to-WM contrast were subsequently quantified (see Methods for details). Although eight breath-holds were averaged in the main results of our work, we also studied the effect of averaging fewer breath-holds to determine whether shorter scan time (fewer breath-holds) would yield sufficient CNR.

Figure 3 shows the subject-averaged CNR at 3T and 7T when modifying the number of breath-holds averaged (Figure 3A and 3B), along with a subject-wise regression for CNR values between 3T and 7T (Figure 3C). CNR and GM-to-WM contrast were significantly higher at 7T relative to 3T (p_CNR_ = 0.0012; p_GM-to-WM_ = 0.0009) (Table 1). As more breath-holds were averaged, CNR increased significantly for both 3T (*p* = 0.00006) and 7T (*p* = 0.00007); at both field strengths, we found that CNR gains fit to a radical (^___^; R^2^ = 0.99) as a function of boluses averaged (Figure 3B). Of note, the subject-wise regression of CNR values between field strengths (Figure 3C), which provides some insight into subject repeatability, was very high (R^2^ = 0.7). According to the regression slope, CNR values were 1.45 times higher at 7T relative to 3T.

**Figure 3.**
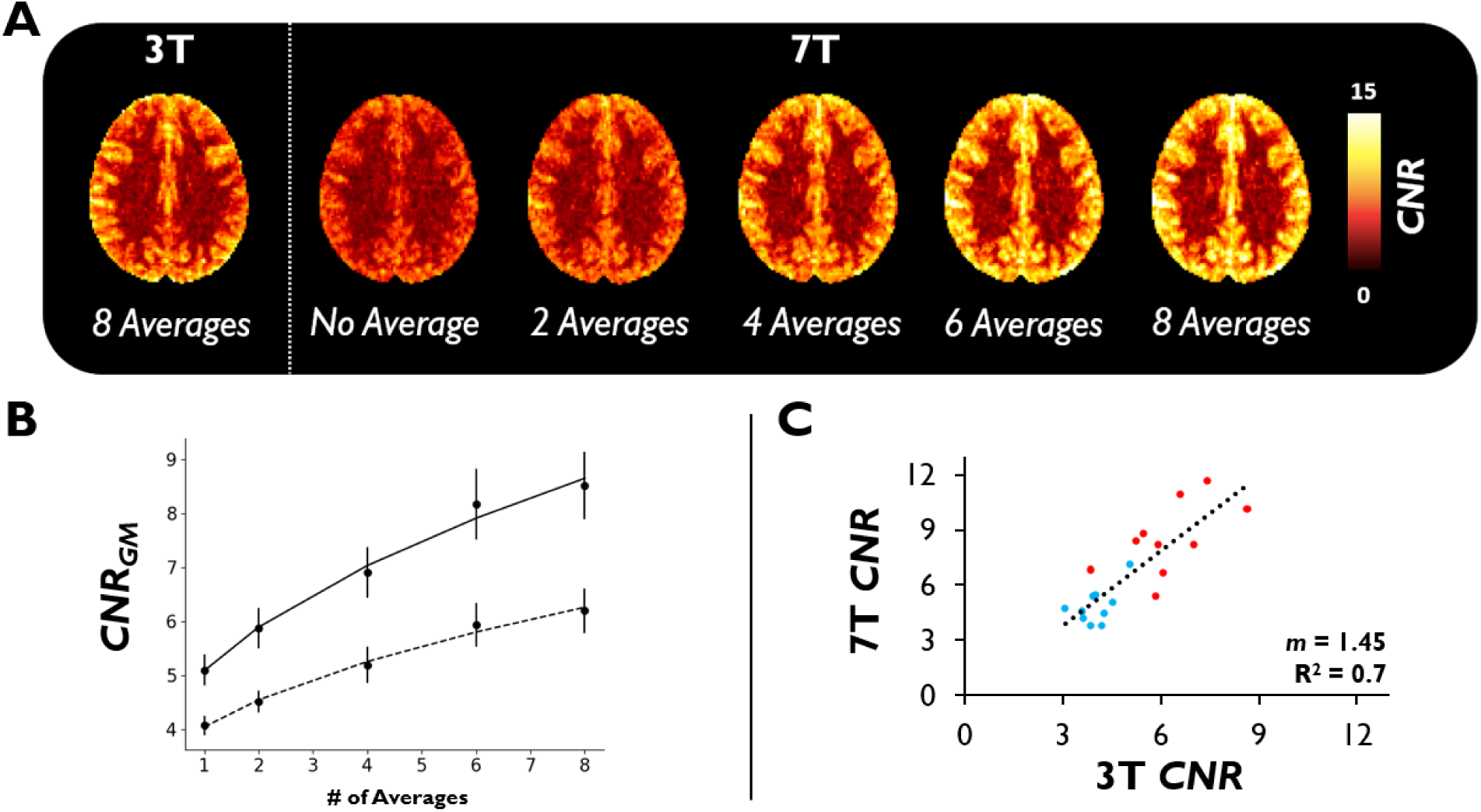
Breath-hold *CNR* Properties. **A.** *CNR* maps were calculated (Eq. 2), transformed to MNI152 2 mm anatomical space, and averaged across subjects for both 3T and 7T (axial view is slightly dorsal to the lateral ventricles). For illustration purposes, *CNR* maps are shown when there is no bolus averaging (1) or 2, 4, 6, or 8 boluses averaged at 7T. **B.** Plot of subject-averaged *CNR* _GM_ at 3T (dashed) and 7T (solid) as a function of the number of breath-holds averaged (error bars represent the standard error). Data are fit to a radical function (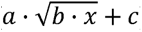) to demonstrate agreement with the well-known relationship between 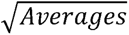 and *CNR*. **C.** Linear regression of subject-wise GM (red) and WM (blue) *CNR* values at 3T vs 7T. m represents the regression slope.

**Table 1.** Summary of bhDSC Perfusion Statistics. Average (mean), standard deviation (std), and coefficient of variation (CV) values for 3T and 7T are shown.

| | | $\Delta S$<br>(%) | | CNR | | CBV<br>(a.u.) | | CBF<br>(a.u.) | | MTT<br>(s) |
| --- | --- | --- | --- | --- | --- | --- | --- | --- | --- | --- |
|  |  | GM | WM | GM | WM | GM | WM | GM | WM | GM |
| 3T | Mean | 1.78 | 1.28 | 6.2 | 4.01 | 5.99 | 4.3 | 59.34 | 43.14 | 6.96 |
|  | Std | 0.32 | 0.13 | 1.31 | 0.55 | 1.16 | 0.97 | 20.69 | 16.13 | 2.11 |
|  | CV | 0.18 | 0.10 | 0.21 | 0.14 | 0.19 | 0.23 | 0.35 | 0.37 | 0.30 |
| 7T | Mean | 2.2 | 1.27 | 8.52 | 4.8 | 5.2 | 3.3 | 44.29 | 25.48 | 7.36 |
|  | Std | 0.44 | 0.43 | 1.96 | 0.99 | 2.15 | 1.39 | 10.05 | 6.05 | 2.52 |
|  | CV | 0.20 | 0.19 | 0.23 | 0.21 | 0.41 | 0.42 | 0.23 | 0.24 | 0.34 |

| | | $\Delta S$<br>(%) | | *CNR* | | *CBV*<br>(a.u.) | | *CBF*<br>(a.u.) | | *MTT*<br>(s) |
| --- | --- | --- | --- | --- | --- | --- | --- | --- | --- | --- |
|  |  | GM | WM | GM | WM | GM | WM | GM | WM | GM |
| 3T | Mean | 1.78 | 1.28 | 6.2 | 4.01 | 5.99 | 4.3 | 59.34 | 43.14 | 6.96 |
|  | Std | 0.32 | 0.13 | 1.31 | 0.55 | 1.16 | 0.97 | 20.69 | 16.13 | 2.11 |
|  | CV | 0.18 | 0.10 | 0.21 | 0.14 | 0.19 | 0.23 | 0.35 | 0.37 | 0.30 |
| 7T | Mean | 2.2 | 1.27 | 8.52 | 4.8 | 5.2 | 3.3 | 44.29 | 25.48 | 7.36 |
|  | Std | 0.44 | 0.43 | 1.96 | 0.99 | 2.15 | 1.39 | 10.05 | 6.05 | 2.52 |
|  | CV | 0.20 | 0.19 | 0.23 | 0.21 | 0.41 | 0.42 | 0.23 | 0.24 | 0.34 |

### Novel Determination of an AIF for bhDSC

A prerequisite for DSC MRI is the presence of signal change upstream of tissue (typically at the arterial level), which is then used as an input function (i.e., AIF) for perfusion quantification in the tissue and veins. In Gd- or hypoxia-based methods, arterial signal change results from an increase in paramagnetic contrast agent in the artery, which subsequently passes through to the tissue.^[6],[8]^ However, given that blood oxygenation in healthy subjects is typically fully saturated in the major cerebral arteries, an increase in blood flow induced by CO_2_ following breath-holding is not expected to significantly change the amount of dOHb in the arteries.^[26],[27],[28]^ Consequently, arterial magnetic susceptibility is not expected to change much as a result of breath-holding. However, hypercapnia uniquely results in vasodilation^[29]^— therefore, while MRI signal is not expected to change in the arteries from paramagnetic contrast agent, we expect that arterial vasodilation will result in a signal decrease, particularly at higher magnetic field strength (see Discussion). Thus, we developed a novel, alternative framework for selecting the AIF when using breath-holds (or hypercapnic paradigms in general).

Figure 4 shows voxels exhibiting a negative signal change (i.e., a positive relaxation change), and, following averaging of these voxels in the middle (MCA), posterior (PCA), and anterior cerebral arteries (ACA) (refer to Methods for selection criteria), displays the subject-averaged AIF time courses at 3T and 7T in response to breath-holding. Voxels with a positive relaxation rate change are located adjacent to the ventricles and in regions containing/adjacent to veins or arteries—the magnitude of the associated relaxation rate changes are significantly higher (p < 0.0007) at 7T in comparison to 3T. The presence of voxels displaying a positive relaxation rate change is, at first glance, counterintuitive given that th predominant effect of breath-holding is a decrease in dOHb (i.e., decrease in the relaxation rate). However, when we simulated (Figure S2) a voxel containing a large vessel, with little-to-no change in dOHb in combination with vasodilation, a substantial relaxation rate increase was in fact observed, in agreement with the experimental results. Thus, the simulations and experimental findings support the hypothesis that arterial vasodilation is responsible for the substantial relaxation rate increase observed in the arteries as a result of breath-holding.

**Figure 4.**
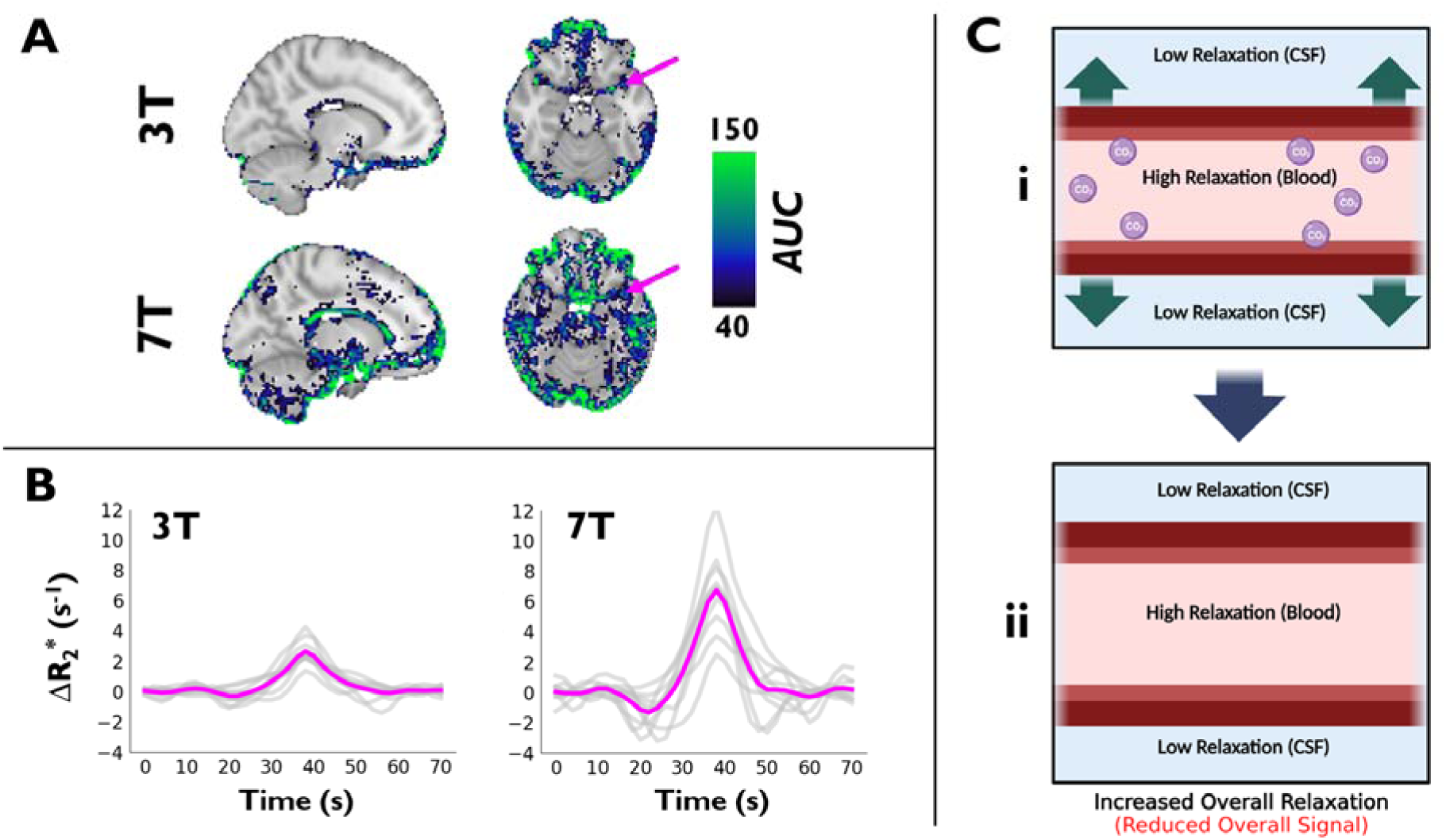
The AIF. **A.** Apparent Δ*R* _2_*\** AUC maps were calculated subject-wise, transformed to MNI152 2 mm anatomical space, and averaged at both 3T and 7T (sagittal and axial views are shown at the level of the middle cerebral artery). Pink arrows indicate an example of the AIF’s location. AUC maps are overlaid onto the MNI152 2 mm anatomical template. **B.** The associated apparent AIF Δ*R* _2_*\** time courses are shown at 3T and 7T, with subject-wise time courses in gray and subject-averaged time courses in pink. **C.** Illustration of vasodilation mechanism, which results in arterial signal decrease during breath-holding.

### bhDSC Perfusion Measurement

The AIF was first scaled by the integral of the VOF, composed of voxels in the superior sagittal sinus (SSS)—this step was conducted as, like in tissue, the VOF integral is representative of the amount of contrast agent (dOHb), whereas, unlike in tissue, the AIF integral is representative of the degree of vasodilation. Following this scaling step, CBV, CBF, and MTT maps were calculated using a standard, truncated singular value decomposition (SVD) analysis, with an SVD noise threshold of 20% and hematocrit correction factor of 1.45.^[11],[30],[31],[32],[33]^ Please note that the reported perfusion values ar considered **relative**, as these values are scaled by some unknown set of factors which depend on acquisition, analysis, and tissue parameters (see Methods)^[11]^; therefore, we report the *CBV* and *CBF* values as (a.u.). As will be discussed, the scaling is not random but fully determined by acquisition and analysis parameters.

Figure 5 shows the subject-averaged *CBV* and *CBF* perfusion maps (Figure 5A), subject-wise GM and WM perfusion values (Figure 5B), and voxel-wise regressions for *CBV* and *CBF* between 3T and 7T (Figure 5C). At 3T, *CBV* values are slightly higher (*p* = 0.108) and *CBF* values are significantly higher (*p* = 0.0027) than at 7T (Figure 5B). The attained GM-to-WM perfusion contrast (i.e., ratio of GM-to-WM perfusion values) was significantly higher at 7T relative to 3T for both *CBF* (*p* = 0.018) and *CBV* (*p* = 0.016) values. In addition, as shown in the subject-wise *CBF* maps (Figure S3), clear GM-to-WM contrast is observed in all subjects at 7T, but not at 3T. High coefficients of determination are associated with the voxel-wise linear regression of 3T and 7T *CBV* (voxel-wise R^2^ = 0.53) and *CBF* (voxel-wise R^2^ = 0.58) values (Figure 5C). This regional agreement is further supported by GM-normalized *CBV* and *CBF* maps (Figure S4); more pronounced differences can be observed in the WM, although these values are not reliable in bDSC or ASL due to lower CNR and longer vascular transit times. Of note, MTT values in GM (Figure S5) are slightly higher, although not significantly (*p* = 0.74), at 7T as opposed to 3T (MTT_7T,GM_ = 7.36 ± 2.52 vs MTT_3T,GM_ = 6.96 ± 2.11). Due to low CNR, MTT values are not reported from the WM.

**Figure 5.**
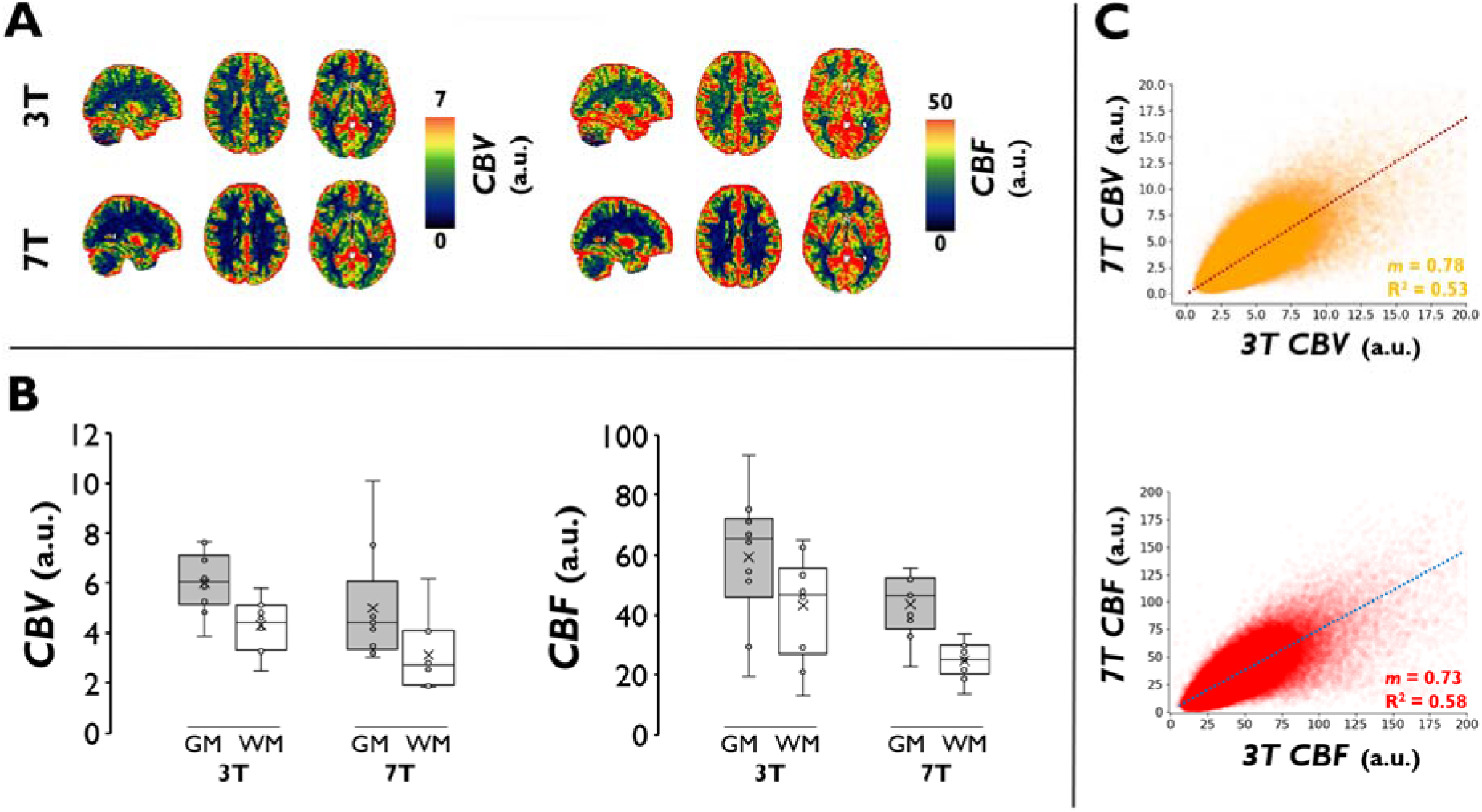
bhDSC Perfusion Results. **A.** *CBV* and *CBF* maps were calculated subject-wise, transformed to MNI152 2 mm anatomical space, and averaged at both 3T and 7T (sagittal (x = 30) and axial views (z = 38, 46) are shown). **B.** Box and whisker plots show the calculated *CBV* and *CBF* values (for GM and WM) at 3T and 7T. C. Voxel-wise regressions for *CBV* and *CBF* at 3T vs 7T. The regressions are based on voxels from the subject-averaged perfusion maps, and only voxels from the GM and WM are displayed. Regressions are set to intersect with the origin. m represents the regression slope.

### Comparing bhDSC with ASL

For validation, we compared our bhDSC results with 3T ASL data obtained from the same subjects (refer to Methods for calculations of *CNR* and *CBF* for ASL). Note that *CBF* values for ASL are not absolute, as a calibration scan was not obtained—thus, only a relative comparison is conducted for *CBF*.

Figure 6 provides CNR (Figures 6A and 6B) and *CBF* (Figures 6C and 6D) comparisons between bhDSC and ASL. bhDSC at 7T yielded significantly higher CNR (*p* = 0.0043) and GM-to-WM contrast (p < 0.0001) in comparison to ASL. At 3T, bhDSC yielded higher CNR (insignificant; *p* = 0.21) but lower GM-to-WM contrast (insignificant; *p* = 0.28) in comparison to ASL. *CBF* maps are regionally congruent between bhDSC and ASL (Figure 6C), supported by the voxel-wise coefficient of determination (voxel-wise R^2^ = 0.51) between 7T bhDSC and ASL *CBF* values (Figure 6D), with largest deviations observed in surrounding vascular territories. Although not shown, the coefficient of determination is lower between 3T bhDSC and ASL (voxel-wise R^2^ = 0.24).

**Figure 6.**
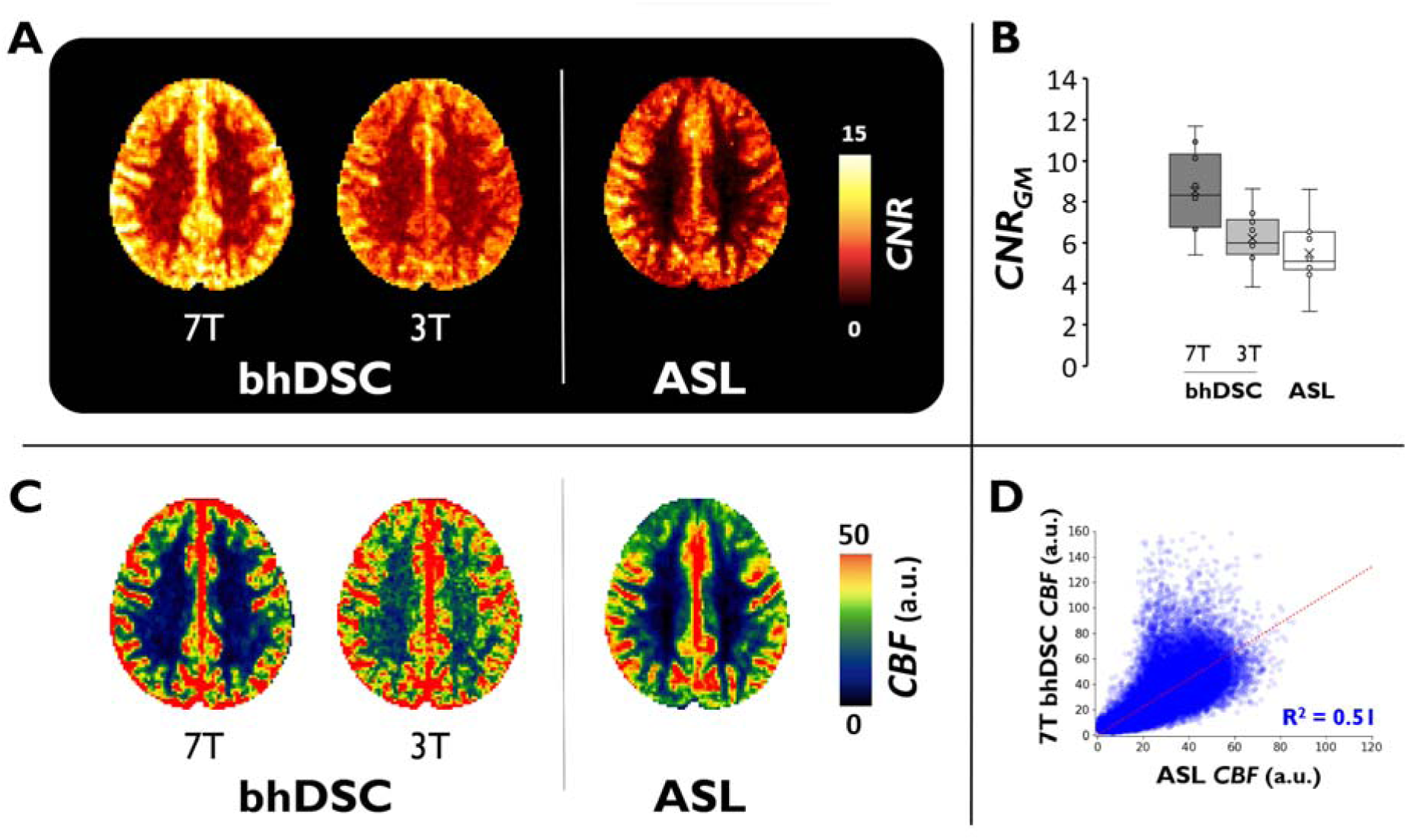
bhDSC vs ASL. **A.** *CNR* maps were calculated (Eq. 2), transformed to MNI152 2 mm anatomical space, and averaged across subjects. **B.** Box and whisker plots show calculated *CNR* values in GM for ASL and bhDSC at 3T and 7T. **C.** *CBF* maps were calculated subject-wise, transformed to MNI152 2 mm anatomical space, and averaged across subjects. Axial view is slightly dorsal to the lateral ventricles for *CNR* and *CBF* maps. **D.** Voxel-wise *CBF* regression between ASL and 7T bhDSC. Regression is based on GM and WM voxels from the subject-averaged perfusion maps. As ASL coverage was variable between subjects, masking was conducted to only include voxels where ASL coverage overlapped amongst all subjects—refer to Figure S6 for mask. We constrained analysis to axial slices 48 through 58, where ASL image intensity was homogeneous across each axial slice. Regression is set to intersect with the origin.

### Effect of Vasodilation on CBV Estimation

Given that bhDSC relies on dynamic vasodilation for bolus generation in the tissue, unlike in other methods such as ASL or Gd-DSC, it is unclear as to whether baseline perfusion measures can truly be estimated during a vasodilatory challenge. To address this concern, we developed simulations (refer to Supplementary Materials)^[11]^ to assess the magnitude by which tissue vasodilation introduces error into calculated baseline perfusion measurements.

Figure 7 shows the effect that simulated tissue vasodilation has on *CBV* quantification. According to the simulations, the induced vasodilation that occurs during a breath-hold results in *CBV* underestimation, relatively independent of baseline *CBV* values and tissue composition. Given an approximate relative vasodilation (ΔCBV) of 4-9% during breath-holding, the associated *CBV* underestimation is ∼20-40%. Thus, for example, a *CBV* of 4% would be underestimated to be around 2.8%. Note that a range is provided as, depending on breath-hold performance, the amount of vasodilation will vary between subjects and, consequently, the underestimation of *CBV* values will be subject dependent. Although globally underestimated, it appears that regardless of tissue blood volume or vessel composition, underestimation does not vary much and, therefore, regional scaling differences are not expected. In other words, even though the absolute quantitative values are affected by vasodilation, the relative distribution of values remains largely unaffected. However, in the event that different voxels have substantial variations in vasodilatory capacity (reflected by more variability in the x-axis for a given subject), it can be expected that regional scaling differences due to vasodilation will be observed (see Discussion). Although not shown here, *CBF* underestimation from vasodilation shows the same behavior as *CBV* underestimation; thus, the *MTT* is not notably affected.

**Figure 7.**
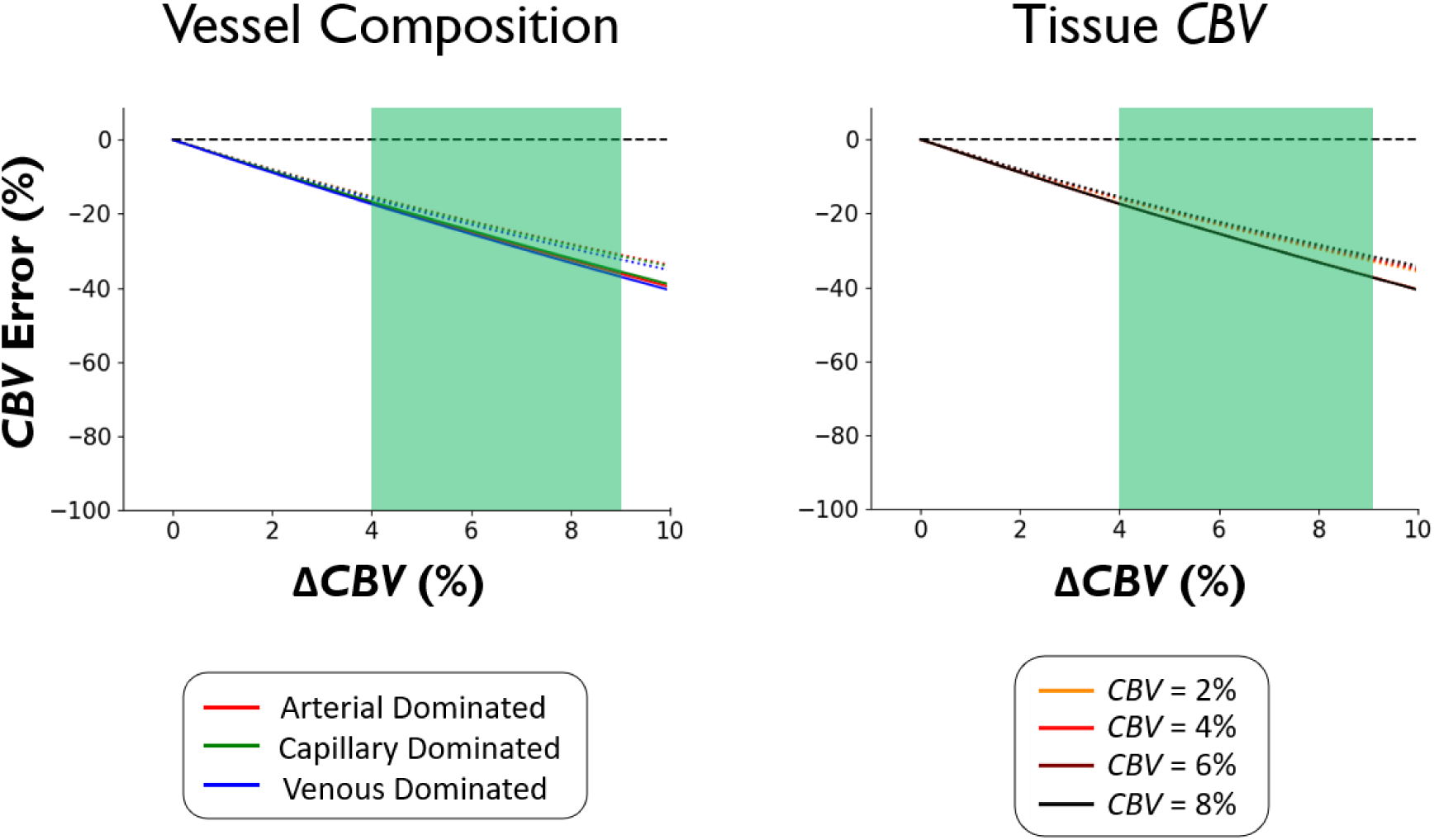
Effect of Vasodilation on *CBV* Quantification. The percentage error in *CBV* quantification as a direct consequence of vasodilation (Δ*CBV*) was simulated by varying the amount of vasodilation that occurs in tissue during a breath-hold. Values on the x-axis are relative, that is, a Δ*CBV* of 5% for a tissue baseline *CBV* of 4% results in a final *CBV* of 4.2%. The green area (4 - 9%) represents the typical range of vasodilation that occurs during a 16 s breath-hold (refer to Simulation Methods). All values below the black dashed line represent an underestimation as a direct consequence of vasodilation. Solid lines represent 7T results whereas dotted lines represent 3T results. **Left**. Results are shown for simulated voxels with varying vessel compositions (described in Table S1). Tissue *CBV* is 4% for this simulation. Right. Results are shown for simulated voxels with varying tissue *CBV*. Voxels are venule dominated for this simulation.

## Discussion

For the first time, we performed a DSC-MRI perfusion analysis using a breath-hold task, without exogenous contrast or additional medical device equipment. We have found the following:

⍰ The AIF during a breath-hold task is uniquely characterized by a negative signal change, likely representing vasodilation, as supported by both simulations and the literature.
⍰ bhDSC-calculated *CBV* and *CBF* are generally within the physiological range of values reported in the literature using established DSC-MRI approaches.
⍰ bhDSC-calculated *CBF* demonstrates regional congruency and voxel-wise linear agreement with *CBF* determined using ASL.
⍰ At 7T, the breath-hold task yielded significantly higher *CNR* (*p* = 0.0012) and GM-to-WM contrast (*p* = 0.0009) relative to 3T and ASL (*p* = 0.0043 and *p* < 0.0001, respectively).
⍰ Tissue vasodilation, unique to bhDSC, yields a global *CBV* and *CBF* underestimation of ∼20-40%, but assuming relatively homogeneous vasodilatory capacity, error is consistent across voxels with different vascular properties.

### From Breath-Hold to Signal Change

It is well established that hypercapnia induced by breath-holding produces a quantifiable T_2_* signal increase in the tissue, which has been particularly useful in CVR and calibrated BOLD studies.^[24],[25],[34],[35]^ Of note, mild hypoxia is also known to accompany hypercapnia during breath-holding, although, given a ∼0.65% reduction in arterial oxygen saturation during a 16 s breath-hold,^[27],[36]^ the impact of breath-hold-induced hypoxia on T_2_* signal change is expected to be negligible in comparison to the effect attributed to hypercapnia.

It has long been known that increased P_a_CO_2_ (arterial partial pressure of CO_2_) leads to increased blood flow in the arteries, and subsequently, in tissue capillaries and veins.^[37]^ This occurs either directly through CO_2_ acting on the smooth muscle cells surrounding the arteries/arterioles, or indirectly through CO_2_ acting on the vascular endothelium, in both cases resulting in vasodilation and increased blood flow.^[21]^ Therefore, a breath-hold, which yields a transient rise and fall (i.e., bolus) of P_a_CO_2_, is a relatively simple way to induce a transient bolus of *CBF*. Of note, the breath-hold duration has a sigmoid-like relationship with the resulting change in P_a_CO_2_,^[36]^ which then also has a sigmoid-like relationship with the resulting change in *CBF* ^[38]^. Given that oxygen metabolism is assumed to be constant during mild hypercapnia,^[26],[28],[23],[39],[40]^ the *CBF* bolus resulting from arterial/arteriolar/capillary vasodilation now permits the cerebral tissue to be supplied with more oxygen than it requires, leading to an increase in blood oxygenation and a reduction in dOHb, in accordance with the bolus shape and magnitude of P_a_CO_2_ in tissue.

We refer to what has just been described as the CVR hypothesis, wherein the observed arterial, tissue, and venous signal changes are a result of CO_2_-induced vasodilation throughout the vascular tree. That is, the increased CO_2_ following breath-holding leads to an increase in arterial blood volume, in turn resulting in increased flow. In this hypothesis, it is predominantly CO_2_-induced vasodilation/flow increases prior to the tissue which result in an oxygenation bolus at the tissue level. However, a second mechanism—the cardiac hypothesis—may alternatively describe the observed oxygenation bolus. Here, the previously described CVR effect may occur in addition to a cardiac output increase, which propagates throughout the cerebral vasculature as a bolus of increased blood flow (in accordance with the breath-hold duration). According to the literature, the likelihood of a cardiac hypothesis is very low given that cardiac output can increase, decrease, or remain constant depending on the breath-hold maneuver.^[41]^ In fact, Sakuma *et al.* found that large lung volume breath-holding (end-inspiration) results in a reduction of cardiac output, which would theoretically yield the reverse effect of that observed in our data.^[42]^

Nevertheless, these mechanisms predict a change in paramagnetic dOHb concentration in the tissue vasculature and draining veins, which is the basis for T_2_* signal change in fMRI.^[43],[44],[45],[46]^ The magnitude of these signal changes scale with the underlying baseline *CBV*, allowing for measurement within a DSC framework. In the current study, we observed a ∼10-15 s delay between the initiation of hypercapnia and the resulting T_2_* signal change bolus onset (Figure 1), which is similarly observed in previous hypercapnia studies and is attributed to the time for CO_2_ build-up in the lungs and the lung-to-brain travel time of blood.^[47],[48]^ Ultimately, the decrease in dOHb results in a transient increase in MRI signal, and the magnitude of the signal change is dependent on the magnitude of ΔP_a_CO_2_, field strength, pulse sequence, and tissue composition.^[49]^

### Determination of an AIF for bhDSC

To measure *CBV*, *CBF,* and *MTT* using DSC MRI, the tissue bolus must be deconvolved with an input bolus, ideally from the cerebral arteries (AIF), that acquires dispersion and delay as it travels to and through the tissue, in accordance with indicator dilution theory.^[6],[8]^ While an AIF is expected and routinely determined when using Gd or hypoxia contrast, the presence of an AIF and its associated properties are unknown in bhDSC. To address this, we examined the physiological processes underlying arterial signal change as a result of breath-holding.

As argued above, dOHb levels in healthy subjects do not change much in the arteries during breath-holding as there is effectively no oxygen exchange at this level of the vasculature and arterial blood oxygenation is already near-saturation.^[26],[28]^ Thus, given that most voxels in a typical T_2_*-weighted MRI acquisition (2-4 mm voxel dimensions) likely consist of more than one tissue/vessel type, any observed signal increase in and around the MCA, for example, may represent signal changes from surrounding, early-perfused cortical tissue and nearby veins, and using this as an AIF would violate basic tenets of indicator dilution theory.

Experimentally, we noticed that the MRI signal, which increased throughout most of the brain in response to a breath-hold, decreased in many voxels containing large arteries and/or veins, particularly when adjacent to CSF, with a larger observed effect at 7T (Figure 4). To better understand this phenomenon, we modeled (Figure S2) an arterial voxel with vasodilation (i.e., *CBV* change) but no dOHb change. In doing so, we found that vasodilation resulted in a sizable signal decrease (i.e., apparent relaxation rate increase) in accordance with our experimental findings. The reason for this phenomenon relies on the fact that the intravascular transverse relaxation rate is substantially higher than the extravascular tissue and CSF relaxation rates at 7T, and to a much lesser extent, at 3T, for which the intravascular transverse relaxation rate is higher than CSF relaxation.^[50]^ Thus, an increase in *CBV* with little-to-no change in blood oxygenation leads to an MRI signal decrease (Figure 4C). This signal origin has also been hypothesized as an explanation for the initial signal dip observed in fMRI studies^[49]^ and has been posited in other work.^[51],[52]^ Although its magnitude depends on vessel orientation and voxel composition, measurable signal change is expected from our simulations to occur in any voxel where a relatively large vessel increases in volume with negligible change in blood oxygenation, displacing extravascular tissue and/or CSF, particularly at 7T. Of note, this means that tissue voxels co-localized with larger vessels and CSF may display **both** an initial signal decrease from vasodilation and a subsequent signal increase from the resulting dOHb decrease.

Our findings are also supported by the literature, where negative signal change during hypercapnia was found in and adjacent to the ventricles.^[47],[53]^ At 3T, this effect results almost entirely from the difference between CSF and intravascular signal, as previously described.^[53]^ Voxels characterized by a signal decrease were observed in higher abundance at the arterial and venous level in a previous fMRI study at 7T,^[54]^ as is similarly observed here.

While we were unable to sample P_a_CO_2_ in the cerebral arteries while directly imaging changes in vascular occupancy—a potential avenue for future work—there is very strong experimental and theoretical evidence that the apparent positive T_2_* relaxation rate change at the arterial level represents a time course of arterial vasodilation in response to changes in P_a_CO_2_. This phenomenon is not expected in Gd or hypoxia studies, where relaxation rate is known to increase in the artery due to increased paramagnetic contrast agent, and not vasodilation. Although it is not the traditional input of contrast agent-induced signal change, arterial vasodilation does in fact represent the input induced by hypercapnia, likely reflecting the input P_a_CO_2_ time course in the brain, which then passes through and acts on downstream vasculature to yield tissue signal change in accordance with the shape and magnitude of the P_a_CO_2_ bolus. The careful selection and averaging of voxels in and around the MCA, PCA, and ACA exhibiting this apparent positive relaxation rate change provided us with a novel solution for deconvolving the tissue data with an input function. The fact that arterial signal change is opposite in sign to signal change observed in much of the brain will likely make it easier to identify an AIF using bhDSC in comparison to traditional DSC techniques. Again, given that we have not acquired P_a_CO_2_ time courses in our data, future work would need to establish the quantitative relationship between P_a_CO_2_ and MRI signal change in and around the arteries.

Of note, we recognize that vasodilation, and thus negative signal change, may also arise at the arterial level from other sources, such as changes elicited in the cardiac cycle. However, we are confident that these extraneous effects are either aliased due to the lower temporal resolution (TR = 2 s) or averaged out during the process of breath-hold bolus averaging.

### CNR in bhDSC

There is a significant increase in *CNR* and GM-to-WM contrast as field strength increases (Figure 3; Table 1). These findings are supported by our simulations (Figure S2) and studies that have demonstrated a higher transverse relaxivity of dOHb as a function of field strength^[49],[55],[56]^. Thus, for the same breath-hold duration and performance, higher signal change will be observed at 7T relative to 3T. In fact, 1 min of scan time at 7T resulted in approximately the same *CNR* and GM-to-WM contrast as ∼4 min of scan time at 3T. This supports the notion that a bhDSC analysis will yield more robust and, hence, clinically valuable results at higher field strength, within a shorter scan duration. Although there is also a significant increase in *CNR* as the number of breath-holds averaged increases, we found the smallest *CNR* and GM-to-WM contrast gains when comparing 6 *vs* 8 breath-holds averaged (Figures 3A and 3B), in agreement with the known *CNR* ∝ 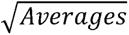 relationship. Our data suggest that bhDSC (TA: 8.5 min) yields superior *CNR* with respect to ASL (TA: 6.5 min), particularly at 7T where *CNR* is ∼1.6 times higher than ASL *CNR*. Of note, bhDSC *CNR* at 7T is even significantly higher than ASL *CNR* (*p* < 0.05) when using 6 breath-holds (TA: ∼6.5 min). As both methods are non-invasive, this is an important finding given that *CNR* is a known limitation of ASL, and bhDSC can yield additional perfusion parameters (*CBV* and *MTT*) in comparison to standard ASL.

The small increase in WM *CNR* as a function of field strength requires closer examination. This finding may be partially explained by the fact that fMRI signal in large pial veins is known to increase more than linearly as a function of field strength (and these veins are often co-localized to the GM mask). Thus, we expect that to some extent, GM *CNR* will increase more than WM *CNR* as a function of field strength, leading to a difference in the GM-to-WM ratio. It is also possible that the calculated GM-to-WM perfusion ratio varies as a function of field strength for reasons other than this partial volume effect, although this is unclear. Until validated further, bhDSC may be most useful for studying GM perfusion and large vessel flow.

### bhDSC Perfusion Measurement

The subject-averaged *CBV* and *CBF* values and maps (Figures 4 and S3; Table 1) are generally within the documented range of previously reported values (*CBV* _GM_ = 3-11 mL/100g; *CBF* _GM_ = 52-137 mL/100g/min).^[30],[57],[58]^ While high GM-to-WM contrast is observed in all individual subject maps at 7T, this is not always the case at 3T (Figure S3), suggesting that bhDSC will yield perfusion maps that are more reliable at higher field strength.

The discrepancy between 3T and 7T values supports the notion that DSC measurement is relative, with a dependence on field strength, amongst many other parameters. The high-level explanation for the *CBV* and *CBF* measurement discrepancy is that tissue *AUC* increases less than VOF *AUC* as field strength increases (Figures 2 and S2). Specifically, we found that *AUC* _GM_ nearly doubles while the average *AUC* _VOF_ nearly triples as field strength increases from 3T to 7T. Given that *CBV* is calculated as a ratio of *AUC* _GM_ to *AUC* _VOF_ (and *CBF* is scaled by this ratio), the aforementioned scaling difference results in lower calculated values of *CBV* and *CBF* at higher field strength—this is also supported by our simulations (Figure S2). This scaling difference likely results from the non-linear intravascular contribution, which is a larger signal component at lower field strength (approximately 10-40% of the signal at 3T in comparison to about 0-5% of the signal at 7T in accordance with Uludag *et al.,* 2009).^[49]^ To reiterate, as there is no commonly accepted pipeline for absolute DSC MRI quantification due to the many quantification dependencies that exist, calculated perfusion values are, to-date, considered relative, and only these relative values are considered in clinical care. Nevertheless, there is a strong voxel-wise linear correlation between 3T and 7T tissue perfusion values (Figure 5C) and visual agreement in the GM (Figure S4), indicating that there are no major regional gray matter scaling differences between field strengths in the GM of healthy subjects. We hope that our documentation of the perfusion estimation differences between 3T and 7T will aid in the development of corrective measures in the future, for absolute quantification.

As previously described, the *CBF* maps are generally well correlated between bhDSC and ASL, with the largest deviations observed in surrounding vascular territories (Figure 6C). The observed incongruence between ASL and bhDSC is expected in the arterial and venous territories surrounding the GM: methods such as bhDSC and Gd-DSC track a contrast agent (dOHb and Gd, respectively), which does not traverse the blood brain barrier in healthy subjects, yielding large signal changes in the capillaries/tissue, supplying arteries, and draining venous vasculature. In methods such as ASL and H_2_O-PET, the tracer is labeled water, which in humans, largely exchanges with the tissue, and therefore only yields high signal change in the capillaries and tissue (low signal change in the arteries/veins). Thus, regional incongruency in territories containing large vessels is expected between bhDSC and ASL.

Although we did not acquire data from multiple sessions per subject at a given field strength, we found that when bootstrapping different bolus combinations to calculate GM *CBV* at 7T (i.e., averaging 4 boluses for a total of 70 bolus combinations), the intraclass correlation (ICC = 0.63) was in the moderate to substantial range (Figure S7), with the lowest precision observed in subjects where the breath-hold bolus shape was not well-defined (subjects 9 and 10). In addition, we observed a strong correlation in subject-wise *CNR* between field strengths (Figure 3C). Both of these findings support the notion that bhDSC, at the very least, is a moderately repeatable method.

While *MTT* _GM_ values (Figure S5; Table 1) are twice as high as values obtained in studies using Gd contrast, they are similar to, if not lower than those reported in studies using hypoxia contrast.^[11],[15],[30],[58]^ Once again, this discrepancy likely results from another known relative DSC quantification dependency, namely: we have previously found longer bolus durations to result in higher calculated *MTT* values.^[11]^ A breath-hold of 16 s induces, on average, a 35 s signal bolus, which is substantially longer than a Gd bolus (but similar in duration to a typical hypoxia bolus), and consequently yields a higher calculated *MTT* due to the inherent limitations of SVD. Although slightly higher at 7T, *MTT* measurements are not statistically different between field strengths. Of note, in accordance with the central volume principle,^[31]^ the higher calculated *MTT* values explain the slight underestimation of *CBF* in comparison to average values reported in the literature.^[30],[57],[58]^

Finally, the delay maps are useful for addressing whether the breath-hold induces a bolus that traverses the brain in a manner dependent on the vascular hierarchy/topography, acquiring delay/dispersion as it progresses. According to our delay maps (Figure S8), the bhDSC method yields delay values across the brain in accordance with the ordering of bolus arrival times reported in the literature when using Gd as a contrast agent^[59]^ and breath-holding for CVR analysis.^[60]^ This provides additional evidence that bhDSC involves a bolus whose delay and dispersion properties are dependent on the location within the cerebral vascular network where tissue is perfused.

### Clinical and Physiological Considerations for bhDSC

There are notable benefits associated with bhDSC in comparison to traditional DSC methods, including the lack of exogenous contrast and/or medical device equipment, and their associated limitations (i.e., patient discomfort, contrast agent extravasation, medical device maintenance, and additional costs).^[13],[61]^ Given that multiple perfusion parameters can be acquired without the necessity of (potentially) scanner-restrictive multi-delay acquisitions, there are also notable benefits with respect to ASL. However, it is important to consider whether there are physiological and/or physical processes, unique to breath-holding, which limit bhDSC’s clinical utility in comparison to available state-of-the-art methods.

In a standard DSC analysis,^[32]^ it is assumed that *CBV* and *CBF* do not change, a condition which is fulfilled during mild hypoxia or Gd. In bhDSC, signal change induced by a dynamic vasodilation time course in tissue may confound the calculated perfusion metrics. To investigate, we simulated (see Supplementary Materials for simulation framework) a 16 s breath-hold in addition to vasodilation in tissue and found global *CBV* and *CBF* underestimation by 20-40% (Figure 7). We found the underestimation to be almost identical in magnitude for voxels with differing vascular compositions and *CBV* values, thus, relative values are expected to be preserved across the brain. On the other hand, given the presence of other global quantification errors in DSC, global scaling underestimation from vasodilation is not expected to hinder the utility of this method, particularly if a global correction factor is introduced. Thus, from our simulations, vasodilation in the tissue is not expected to confound the relative distribution of *CBV* values in healthy subjects. However, this may not be the case in patients, for which there are more regional variations in vasodilatory capacity compared to healthy subjects. Here, it may be useful to develop and employ correction strategies for the relative perfusion values in diseased tissue when using bhDSC.

Another consideration is the potential variability in the oxygen metabolism rate (CMRO_2_) during breath-holding, as this would additionally violate an important DSC analysis assumption. There is very compelling evidence that shows a lack of significant CMRO_2_ variability during mild hypercapnia or breath-holding, especially for the short, 16 s breath-hold duration employed in our study.^[26],[28],[23],[39],[40]^

Causality of flow in the vascular network is another important consideration. In healthy subjects, the cerebral vasculature is assumed to respond relatively homogeneously to a vasodilatory stimulus.^[38],[62]^ However, in certain patients (i.e., those with steno-occlusive disease and/or vascular steal physiology), the affected arteries may have reduced vasodilatory capacity, which would result in tissue downstream from the affected vessels yielding reduced calculated *CBV* values in comparison to tissue supplied by other vessels (and potentially negative values in the case of vascular steal)^[63]^, even though the ground truth blood volume values may not be reduced. In these patients, apparent hypoperfusion relative to the rest of the brain may be a result of combined hypoperfusion and reduced vasodilatory capacity in the supplying vasculature, limiting bhDSC’s validity in this cohort without any additional data or correction algorithms.

Finally, unlike Gd- and hypoxia-based methods, the signal change in bhDSC reflects an oxygenation bolus that only begins at the tissue level—thus, signal change in bhDSC is capillary/venous dominated. When comparing bhDSC with Gd-DSC perfusion results, it is expected that there will be some incongruency in brain regions which have substantially more arterial blood. However, this difference also creates an opportunity to combine information from multiple methods to isolate different blood pools.

### Limitations and Future Directions

Given that dOHb has a far lower molar susceptibility than Gd^[11]^ and ΔdOHb is relatively small during breath-holding^[26],[36]^, it can be expected that bhDSC will yield substantially lower *CNR* than Gd-DSC. However, as illustrated in this work, both the accuracy and precision of perfusion measurement with bhDSC is still sufficient on an individual subject level at 7T and on a group level at 3T. At lower field strengths (such as 3T), incorporating physiological and physical noise correction methods and acquiring multi-echo T_2_* acquisitions^[64]^, may enable bhDSC to yield more robust perfusion metrics for individual subjects.

The breath-hold itself confers some limitations in comparison to ASL. Patient compliance is required during the repeated breath-holds of 16 s, which may be difficult for certain subjects, particularly the elderly, and those with neurological or lung disease.^[65],[66]^ However, breath-holds are often applied in patients for body imaging, and as previously described, sufficient contrast can be obtained at 7T with fewer breath-holds, thereby, reducing the duration required for patient compliance. Another limitation is that movement associated with breath-holding may result in detectable motion during the scan. However, in the present study, we found this to be minimal in effect and correctable using post-processing (motion correction). As well, the breath-hold paradigm requires either a projector setup in the scan room (and the associated programming), or auditory cues through headphones. Finally, there is inter-subject variability in the breath-holds, which is reflected in the variability of bolus shape and magnitude between subjects (Figure S1). The variability (i.e., coefficient of variation (CV)) of our calculated perfusion metrics (Table 1) is on par with that observed in a previous study^[15]^ which used a gas control system for DSC perfusion measurement at 3T (*CBV* _CV_ = 0.33; *C B F*_CV_ = 0.42), limiting the concern that the contrast bolus needs to be standardized across subjects to ensure precision of the calculated perfusion values. Nevertheless, precision should be enhanced in future work by optimizing the breath-hold approach—providing more detailed instructions to the subjects, including whether an end-inspiration or end-expiration is employed and, potentially utilizing longer rest durations between breath-holds.^[67]^

While we have performed a direct comparison with ASL, a direct subject-wise bhDSC *vs* Gd-DSC comparison will allow for a stronger validation of the technique’s utility and limitations in perfusion imaging. As well, the clinical utility of the bhDSC technique should be validated in different patient cohorts, to determine whether all or a subset of those with vascular brain pathology will benefit from bhDSC alone or in combination with additional perfusion imaging techniques (i.e., ASL). Finally, to provide more clinical centers with access to this method, it will be prudent to assess the viability of and optimize the bhDSC technique at lower field strengths.

## Conclusion

For the first time, we leveraged breath-holds to measure perfusion with a DSC analysis. This was made possible by developing and applying a novel arterial input function strategy, tailored for breath-hold and hypercapnic paradigms. In doing so, we found that perfusion values (i.e., *CBF* and *CBV*) yielded higher *CNR* and GM-to-WM contrast at 7T, were within the range of physiological and literature-reported values, and demonstrated high regional correlation with ASL *CBF* data obtained on the same subjects. While bhDSC will need to be further validated with other perfusion techniques in healthy subjects and in various diseases, we are convinced that our findings will aid the implementation of contrast-free perfusion imaging, in both basic and clinical research.

## Methods

### Subjects

This study was approved by the Research Ethics Board of Sungkyunkwan University and all procedures followed the principles expressed in the Declaration of Helsinki. Informed consent was obtained in all 10 healthy volunteers (age: 30.4 ± 9.4 years, 3 female). Please note that the same subjects were scanned at both 3T and 7T.

### MRI Sequences and Experimental Protocols

MRI data were acquired at the Center for Neuroscience Imaging Research at Sungkyunkwan University on the Siemens 3T Prisma (Siemens Healthineers, Erlangen, Germany) and Siemens 7T Terra (Siemens Healthineers, Erlangen, Germany) using the commercially available 64 channel head/neck and 32 channel head coils (Nova Medical, Wilmington, USA), respectively.

#### Structural MRI

The 3T parameters were as follows: 3D-MP2RAGE^[68]^ with 1.0 mm isotropic spatial resolution (176 sagittal slices; GRAPPA = 3; Ref lines PE = 32; FoVread = 250 mm; phase-encoding = A-P; TI1/TI2 = 700/2500 ms; L1/L2 = 4°/5°; TE/TR = 2.98/5000 ms; bandwidth = 240 Hz/px; echo-spacing = 7.1 ms; TA = 8:22 min).

The 7T parameters were as follows: 3D-MP2RAGE^[68]^ with 0.7 mm isotropic spatial resolution (240 sagittal slices; GRAPPA = 3; Ref lines PE = 36; FoVread = 224 mm; phase-encoding = A-P; TI1/TI2 = 1000/3200 ms; L1/L2 = 4°/4°; TE/TR = 2.29/4500 ms; partial-Fourierslice = 6/8; bandwidth = 200 Hz/px; echo-spacing = 7.3 ms; TA = 9:15 min).

#### ASL MRI

ASL scans were only acquired at 3T. The ASL data were acquired with 2.0 mm isotropic spatial resolution, FAIR QUIPSS II labeling scheme and 3D gradient-and-spin-echo (GRASE) readout^[69]^ (40 axial slices, FOV = 256 mm x 256 mm, TE/TR = 20.5/4000 ms, TI1/TI2 = 800/1800 ms, bandwidth = 1860 Hz/pixel, EPI factor = 21, segments = 12, turbo factor = 20, no partial Fourier). Four tag-control pairs were acquired for averaging (TA = 6:28 min).

#### Breath-hold MRI

The 3T parameters were as follows: Gradient-echo 2D-EPI (GRE-EPI) with 2.0 mm isotropic spatial resolution (64 interleaved axial slices; GRAPPA = 2; Ref lines PE = 46; SMS = 2; Ref. scan = EPI; FatSat = True; FoVread = 192 mm; phase-encoding = P-A; TE = 30 ms; L = 70°; TR = 2000 ms; bandwidth = 2312 Hz/px; echo-spacing = 0.53 ms; EPI factor = 94) was acquired for the breath-hold experiment (260 TRs, TA = 8:53 min). 6 measurements of opposite phase-encoded (A-P) data were also acquired for distortion correction.

The 7T parameters were as follows: GRE-EPI with 2.0 mm isotropic spatial resolution (74 interleaved axial slices; GRAPPA = 3; Ref lines PE = 54; SMS = 2; Ref. scan = EPI; partial-Fourierphase = 6/8; FatSat = True; FoVread = 192 mm; phase-encoding = P-A; TE = 20 ms; L = 50°; TR = 2000 ms; bandwidth = 2368 Hz/px; echo-spacing = 0.53 ms; EPI factor = 96) was acquired for the breath-hold experiment (250 TRs, TA = 8:46 min). 6 measurements of opposite phase-encoded (A-P) data were also acquired for distortion correction. Please note that the spatial resolution has not been optimized for 7T in this study. We have instead chosen the same spatial resolution as at 3T to allow for easier and direct comparison.

For the breath-hold scans, subjects were instructed to fixate on a black cross at the center of an iso-luminant gray screen and a countdown timer, which indicated when to breathe regularly and when to perform a breath-hold. Specifically, each breath-hold block was composed of three sections: 10 s of preparation with regular breathing, 16 s of breath-holding, and 34 s of regular breathing (Figure 1). This was repeated nine times to obtain multiple boluses for averaging, with an added final baseline of 80 s. Only the first eight boluses were used for each subject as the response to the final breath-hold bolus was not fully included during the acquisition for some subjects.

#### Preprocessing

FSL (version 6.0.4), AFNI (version 23.0.07), and in-house Python scripts (https://github.com/JSchul1998/bhDSC_Scripts) were used for image pre-processing.^[70],[71]^ Anatomical, ASL, and breath-hold T_2_* data from both scanners were corrected for gradient nonlinearities using the Human Connectome Project’s version of the gradunwarp tool (https://github.com/Washington-University/gradunwarp).

The MP2RAGE structural data were pre-processed using presurfer (https://github.com/srikash/presurfer) and skull-stripped using SynthStrip.^[72]^ The background denoised T1-weighted UNI image was segmented using FSL FAST^[73]^ to obtain three tissue classes corresponding to gray matter (GM), white matter (WM), and cerebrospinal fluid (CSF). The breath-hold data underwent slice-timing (3dTshift, AFNI), motion (mcflirt, FSL), and distortion correction (topup, FSL).^[74],[75]^ The pre-processed breath-hold data were linearly detrended and temporally filtered by averaging each signal time point with a 1×5 Gaussian kernel. The first eight breath-hold blocks were then averaged, wherein each of the eight “boluses” was allotted a 72 s window centered at the bolus maximum and averaged—an example of an original and averaged time course is shown in Figure 1. The percentage maximal signal change (*ΔS*) and *ΔS* -to-noise ratio, hereafter termed contrast-to-noise ratio (*CNR*), were then calculated from the bolus-averaged signal time course:

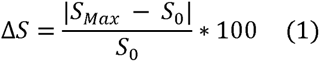

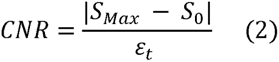

In Eq. 1, *S* _0_ is the average signal of ten temporal volumes before and after (defined as the time that the signal returns to the average pre-bolus baseline) the bolus and *S* _Max_ is the maximum signal increase. The *CNR* is computed as shown in Eq. 2, where ε_t_ is the standard deviation of signal for ten total temporal volumes before and after the selected bolus. GM-to-WM contrast (*GW* _contrast_) was also calculated as follows:

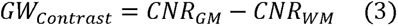

Where *CNR* _GM_ and *CNR* _WM_ represent the mean GM and WM *CNRs*, respectively. *S(t)* was then converted to the change in transverse relaxation rate time course (Δ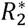(*t*)):

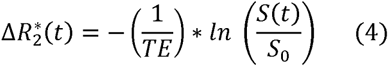

The above equations were applied to T_2_*-weighted breath-hold data from all subjects at 3T and 7T to generate Δ*S*, *CNR*, and 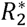(*t*) maps. Note that changes in intravascular or extravascular volume/occupancy, during vasodilation for example, will result in a change in the ‘apparent’ 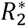(*t*).

The ASL data first underwent motion correction (mcflirt, FSL), resampling to anatomical space (nearest neighbour), and masking (using the skull-stripped anatomical mask).^[75]^ Pairwise subtraction across the four tag-control pairs and subsequent averaging was performed to calculate *CBF* maps. Given the lack of a calibration scan, the *CBF* maps are not absolute, and only a comparison of relative *CBF* maps was conducted between ASL and bhDSC. ASL *CNR* maps were also calculated based on Eq. 2, where *S* _Max_ is the average label signal, *S* _0_ is the average control signal, and ε_t_ is the standard deviation across the controls.

To facilitate group analyses, these maps, along with the later described bhDSC perfusion maps, were transformed to FSL’s MNI152 2 mm space. The processed bhDSC data were first registered to the subject’s anatomical space (6 dof, FSL flirt) and the anatomical data were then non-linearly registered (fnirt, FSL) to the FSL MNI152 2 mm template.^[71],[76]^ The two transformation matrices were then combined into a subject-specific native-to-MNI warp and applied to all subject-specific ASL and T_2_* maps. The MNI space-transformed maps were then averaged across all subjects to generate mean bhDSC_3T_, bhDSC_7T_, and ASL maps. Data were also visually inspected for quality control following each processing step.

### Gray and White Matter Segmentation

GM and WM masks were generated, at both 3T and 7T, using whole-brain anatomical data (FSL FAST). The partial volume estimate maps were then thresholded at 0.9, binarized, and transformed from anatomical to GRE-EPI space for each subject (nearest-neighbor interpolation). Arterial and venous voxels were removed from the masks by thresholding the GRE-EPI temporal standard deviation map (only voxels with values in the lowest 10% of the ε_t_ range were kept in the mask) and using the output to mask the binarized FAST segmentations—the assumption being that voxels with high *CBV* have higher physiological noise due to pulsatility, leading to higher values of ε_t_.^[11],[77]^ The whole-brain GM and WM masks were then used to obtain average GM and WM values (Table 1) from each subject’s ASL, bhDSC perfusion, Δ*S*, and *CNR* maps. GM and WM masks were also generated in standard space from the FSL MNI152 T1 2 mm template using the procedure described above.

### Defining the AIF and VOF

In DSC MRI, it is critical to define an input function that is then used to normalize and deconvolve the tissue data—this input is defined at the level of the artery and is consequently termed the arterial input function (AIF).^[61]^ Typically, to represent the AIF, arterial voxels are identified with Δ*S* values above the 95^th^ percentile and zero delay (see delay measurement below). However, given the spatial resolution of GRE-EPI (typically 2-4 mm in each dimension), signal change derived at the arterial level may be contaminated by signal changes from cortical tissue and nearby veins. With this in mind, we developed an alternative, novel method to obtain an AIF from the breath-hold data, which can be used in any study that uses hypercapnia to calculate perfusion at high or ultra-high field strength. In essence, the selection criteria are the same as those described above, but voxels yielding **negative** *Δ* values, with magnitude above the 99^th^ percentile (*S* _Max_ is replaced with *S* _Min_ in Eq. 1), were used instead. We averaged 15-20 arterial voxels from the middle (MCA), posterior (PCA), and anterior cerebral arteries (ACA). We hypothesize that the resulting signal time course reflects vasodilation (given the negative signal change) at the arterial level (given the short delay and anatomical localization) in response to the CO_2_ stimulus (see Results and Discussion for a more elaborate discussion of the AIF).

In DSC MRI, it is also useful to define an output function from the cerebral draining veins, whose magnitude can be used to scale the measured AIF.^[61]^ Using both GRE-EPI and GRE-EPI-registered anatomical data, 15-20 venous voxels within or adjacent to the superior sagittal sinus (SSS), with Δ*S* values above the 99.9^th^ percentile, long delay (∼ 2-3 s), and low *S* _0_ (as vein contains more dOHb than artery, reducing the baseline GRE signal) were defined and averaged. This averaged venous time course is hereafter termed the venous output function (VOF).

### Perfusion Measurement

All voxel time courses were truncated to commence at the start of the AIF bolus and finish at the end of the VOF bolus. The AIF was then scaled by the integral of the VOF using the following equation:

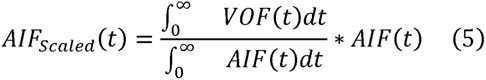

This step was conducted as, like in tissue, the VOF integral is representative of the amount of contrast agent (dOHb), whereas unlike in tissue, the AIF integral is representative of the degree of vasodilation. Normalizing to venous signal also reduces the effect of various perfusion quantification dependencies, such as the baseline oxygenation and susceptibility change induced by the contrast agent.^[11]^

Cerebral blood volume (*CBV*), cerebral blood flow (*CBF)*, and mean transit time (*MTT)* maps were then calculated using a standard, truncated singular value decomposition (SVD) analysis, with an SVD noise threshold of 20%, brain density factor of 1.05, and hematocrit correction factor of 1.45.^[11],[30],[31],[32],[33]^ Although we employ an analysis based on a standard tracer kinetic model, it might be ideal for future work to incorporate Δ*CBF* into the analysis, particularly for the accurate estimation of baseline *CBF* and *MTT*. Note that the perfusion values are **relative**, not absolute; research has shown perfusion imaging values using DSC to depend on various contrast agent and acquisition parameters.^[10],[11]^ Thus, we report values in arbitrary units. Also, note that all voxels yielding a negative *CBV* or *CBF* were removed from subsequent analysis, and are not included in the summary results (Table 1).

### Delay Measurement

Average bolus delay maps were also generated from the bhDSC data at 3T and 7T. Unlike the other perfusion metrics where DSC measurements were conducted in native space, the delay maps were calculated using the group average data.

First, the relaxation time courses were transformed to MNI152 2 mm anatomical space (using the previously described warp matrices) and averaged across subjects. The average time courses were then linearly interpolated to a temporal resolution of 0.5 s. The AIF time courses from each subject were then averaged. The subject-averaged AIF was then shifted forward in 0.5 s steps, up to a maximum of 8 s, and the correlation between the AIF and tissue time courses for each voxel at each temporal shift was recorded. The temporal shift resulting in the highest correlation was determined to be the delay in a model-independent manner. A similar delay method has been employed in previous studies using resting-state fMRI data.^[78],[79]^

Note that the calculated delay maps reflect a combination of artery-to-tissue delay, tissue transit time, and dispersion, given that the AIF time courses were simply shifted to find the maximal correlation with each voxel time course (i.e., there was no separately modeled dispersion parameter in this analysis). The masks used for delay measurement in the occipital GM and putamen come from the MNI structural atlas.

### Statistics

Parameters measured in native space have a calculated mean, standard deviation, and coefficient of variation (CV) for GM and WM, at both field strengths (Table 1). Note that CV is calculated as the inter-subject standard deviation divided by the parameter mean. Linear regressions were conducted for voxel-wise comparisons of *CBV, CBF,* and *MTT*, and subject-wise comparisons of *CNR*. Intraclass correlation (ICC), which measures the proportion of within-subject variance to total variance, was also performed for *CBV*.

We found the data to be normally distributed for all measured parameters in GM and WM using the Shapiro-Wilk test. Thereafter, any statistical testing that was conducted for the experimental data (i.e., comparing mean GM and WM values at different field strengths) was done by applying a two-tailed paired student’s t-test, with an alpha of 0.05.

## Data and Code Availability

The dataset used for the current study is available from the corresponding authors upon reasonable request.

Analysis and simulation code can be found at the following GitHub link:

https://github.com/JSchul1998/bhDSC_Scripts

## Author Contributions

**J.B.S:** Conceptualization, Data Curation, Formal Analysis, Investigation, Methodology, Software, Visualization, and Writing

**S.K:** Data Curation, Software, and Writing

**S.G.K:** Funding Acquisition, Resources, Supervision, and Writing

**K.U:** Conceptualization, Formal Analysis, Funding Acquisition, Investigation, Methodology, Project Administration, Resources, Supervision, and Writing

## Acknowledgments

We would like to thank Boohee Choi and Suji Jeong at the Institute for Basic Science for helping us acquire the data. Figures 1A and 4C were created with BioRender.com.

