## Supplementary Materials for "Non-Invasive Perfusion MR Imaging of the Human Brain via Breath-Holding"

**Simulation Methods**

***Signal Model Overview***

The simulation framework in this study is adapted from a previously developed fMRI/DSC signal model—thus, please refer to these studies for a more detailed discussion of the modeling approach and signal parameters.^[11],[49],[80]^ In essence, the previous model summates signal from extravascular (*EES*) and intravascular (*IVS*) proton spin contributions for simulated arterial, venous, and cerebral tissue voxels at 3T. These signal changes result from dynamic changes in paramagnetic dOHb. As the current study focuses on breath-hold-induced dOHb changes at both 3T and 7T, a few notable modifications were applied, and will now be described in some detail. The signal framework is illustrated in Figure S9.

***Breath-Hold Stimulus***

To simulate the breath-hold (hypercapnic) stimulus in contrast to the previously modeled hypoxic stimulus,^[11]^ some fundamental differences must be considered.

Firstly, as CO_2_ levels increase during a short duration breath-hold, tissue blood oxygenation increases and dOHb concentration decreases—this is the opposite of what occurs during a hypoxic stimulus.^[13],[26],[37]^ An added consideration here is that OHb can only increase to a certain extent depending on the vessel (i.e., assuming artery has a baseline blood oxygenation ($Y_{0}$) of 0.98, oxygenation can only increase by a maximum of 2%; whereas in vein, assuming $Y_{0}$ = 0.65, the oxygenation increase can be much larger). These vascular-dependent cut-offs are incorporated into the modeling of the oxygenation change stimulus ($\Delta Y$). To roughly follow the shape of the experimental breath-hold signal curves, a gamma-variate function was used for the simulation of $\Delta Y(t)$:

$$\Delta Y(t)=a*\left( \frac{t}{b} \right)^{c}*e^{\left( c*\left( 1-\frac{t}{b} \right) \right)} (6)$$

Of note, $a$, which corresponds to the maximum $\Delta Y$ in a given vessel, was informed by previous studies to be approximately 7.5% in vein/venule for a 16 s breath-hold.^[26],[36]^ In artery, $a$ is assumed to be much lower (0.6-1%) given that there is effectively no oxygen exchange at this point in the vascular network.^[26],[16]^ $a$ for capillary was then linearly interpolated and scaled for dispersion to be roughly 5.3%. The other shape parameters for Eq. 6 are included in Table S1.

Another important consideration with respect to a breath-hold is that the hypercapnic stimulus results in vasodilation.^[21],[37]^ The associated vasodilation occurs dynamically and should be included in the modeling in addition to $\Delta Y$, a consideration that is not required for mild-to-moderate hyperoxia-, hypoxia-, or Gd-DSC. For simplicity, the simulated time course of vasodilation ($\Delta$*CBV*(*t*)) roughly follows a gamma variate shape (like that of $\Delta Y(t)$):

$$\Delta CBV(t)=d*\left( \frac{t}{b} \right)^{c}*e^{\left( c*\left( 1-\frac{t}{b} \right) \right)} (7)$$

Note that $\Delta CBV$ is relative here, that is, if $\Delta CBV$ = 0.05 (5%) in a tissue voxel where *CBV* = 4%, the *CBV* would increase to 4.2%. The term $d$, which corresponds to the maximal $\Delta CBV$ in a voxel during breath-holding, was simulated for a range of values, but is likely around 0.04-0.9 (4-9%) for tissue during a 16 s breath-hold. This was derived by using the Grubb’s relationship,^[29]^ Grubb’s exponents of roughly 0.18 for deoxygenated vessels^[81]^—such as capillary, venule, and vein—and 0.29^[82]^—used for arterioles and arteries—during hypercapnia, and a known *CBF* change of around 20-46% for a 16 second breath-hold.^[28],[36],[23],[83],[84],[85],[86]^ The other shape parameters for Eq. 7 are the same as those used for Eq. 6 (included in Table S1). In addition to *CBF* changes from autoregulation, global *CBF* may also change during breath-holding due to changes in cardiac output. For short duration end-exhalation breath-holds,^[42]^ cardiac output has been found to be relatively constant—at most, cardiac output has been shown to decrease during end-inspiration breath-holding.^[41],[42]^

To incorporate dispersion, $\Delta Y\left( t \right)$ and $\Delta CBV(t)$ were convolved with bi-exponential residue functions to obtain the tissue and venous time courses (described in Schulman *et al.,* 2023). The resulting tissue and venous time courses were finally scaled to ensure that the time course maximums corresponded to $a$ and $d$ for $\Delta Y(t)$ and $\Delta CBV(t)$, respectively (Table S1).

Ultimately, it is important to recognize that breath-hold-induced T_2_* signal changes result from an interplay of dynamic blood volume and dOHb changes, dependent on the vascular components of the simulated voxel.

***Extension of Model to 7T***

To extend the previous model to higher field strength, the intravascular and extravascular transverse relaxation rates at 7T were utilized.^[49]^ $R_{2,0,EES}^{*}$ and $R_{2,0,IVS}^{*}$ are the baseline extravascular and intravascular magnetic relaxation rates ($R_{2,0}^{*})$, respectively. At 7T, $R_{2,0,EES}^{*}$ and $R_{2,0,IVS}^{*}$ were experimentally approximated to be 32.6 s^-1^ and 116 s^-1^ for fully oxygenated blood, respectively.^[49]^ $R_{2,con,EES}^{*}$ and $R_{2,con,IVS}^{*}$ are the extravascular and intravascular magnetic relaxation rates induced by dOHb, respectively. While $R_{2,con,EES}^{*}$ at 7T can be determined by referring to Eqs. 4-5 in Schulman *et al.,* 2023, $R_{2,con,IVS}^{*}$ is not readily available and requires extrapolation from $R_{2,con,IVS}^{*}$ at lower field strengths^[49]^—although an approximation, we derived the following equation for 7T:

$$R_{2,con,IVS}^{*}=549*\left( 1-Y_{0}-\Delta Y \right)^{2} (8)$$

Due to its large baseline relaxation, the intravascular relaxation rate change has a minor contribution to the DSC signal at 7T.

***Incorporating a CSF Component***

In this study, we opted to include a CSF signal component ($S_{CSF})$ in the simulated arterial and venous voxels, given that the MCA and SSS are often adjacent to or surrounded by CSF in the subarachnoid space.

$$S_{CSF}=e^{-TE*(R_{2,0,CSF}^{*}+R_{2,con,CSF}^{*})} (9)$$

$R_{2,0,CSF}^{*}$ is the baseline CSF relaxation rate without contrast agent—at 3T, this was measured to be 13.35 s^-1^.^[87]^ At 7T, this value has not been widely reported, but was estimated to be approximately 4.35 s^-1^ from our T_2_* data. This estimation was performed by setting up a T_2_* signal equation^[88]^ at 3T and 7T. The measured CSF signal at each field strength (measured from our T_2_* data) and the literature value of $R_{2,0,CSF}^{*}$ at 3T were then input to the equation to solve for $R_{2,0,CSF}^{*}$ at 7T. $R_{2,con,CSF}^{*}$ is the CSF relaxation rate induced by dOHb—while this was assumed to be similar to the relaxation response in the extravascular space at 3T and 7T, there is likely to be some deviation due to differences in proton density, water diffusion, and molecular composition. Nevertheless, modifying either $R_{2,con,CSF}^{*}$ or $R_{2,0,CSF}^{*}$ within a reasonable range had little effect on the simulated results. CSF signal was then scaled by its volume fraction within the voxel and added to the total voxel signal (Figure S9). The CSF properties are further illustrated in Table S1.

**Supplementary Figures**

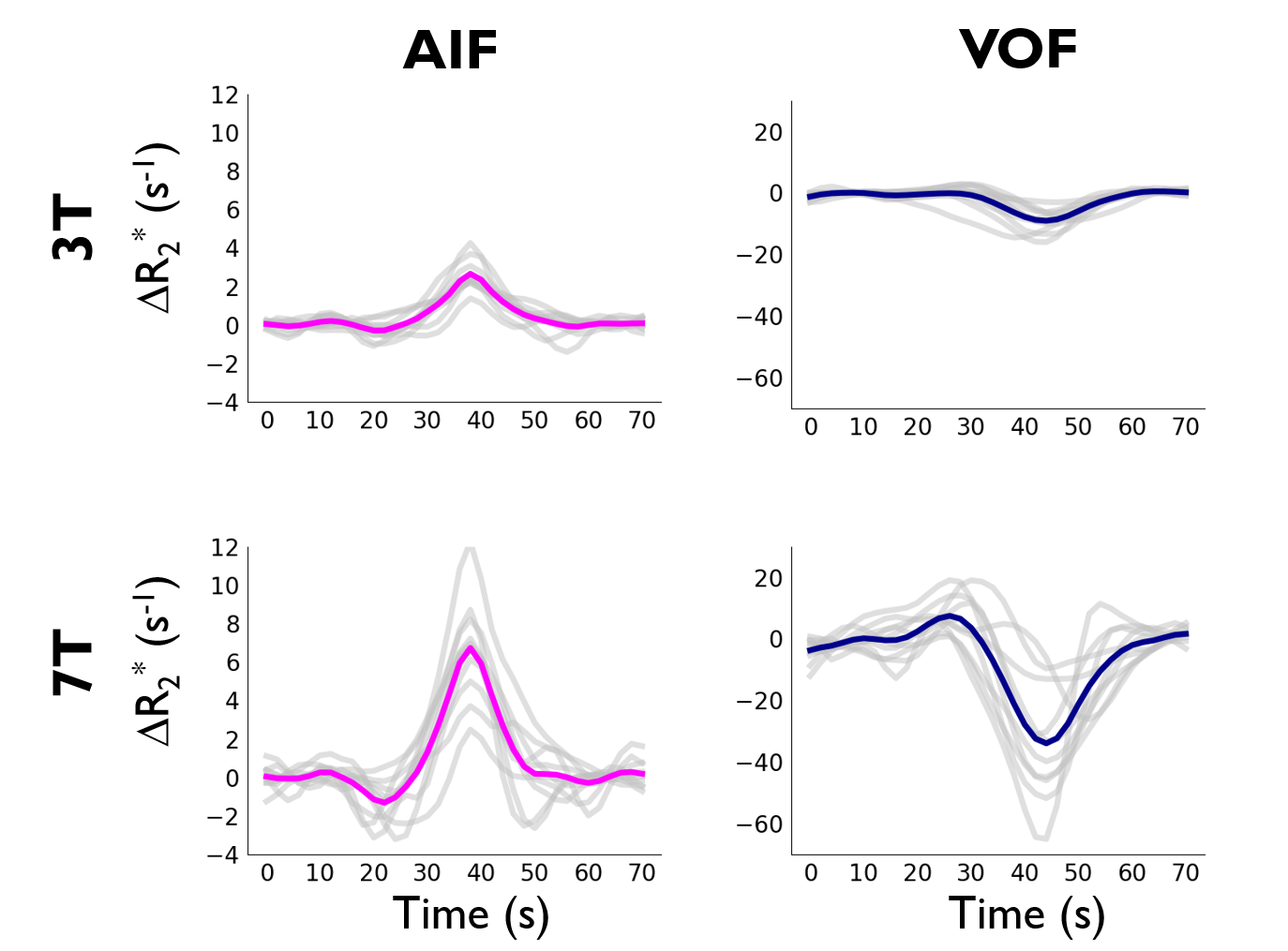

**Figure S1. Subject-wise and Averaged Relaxation Time Courses at 3T and 7T.** Subject-wise time courses are in gray, and averaged time courses are shown in pink for the AIF and navy for the VOF.

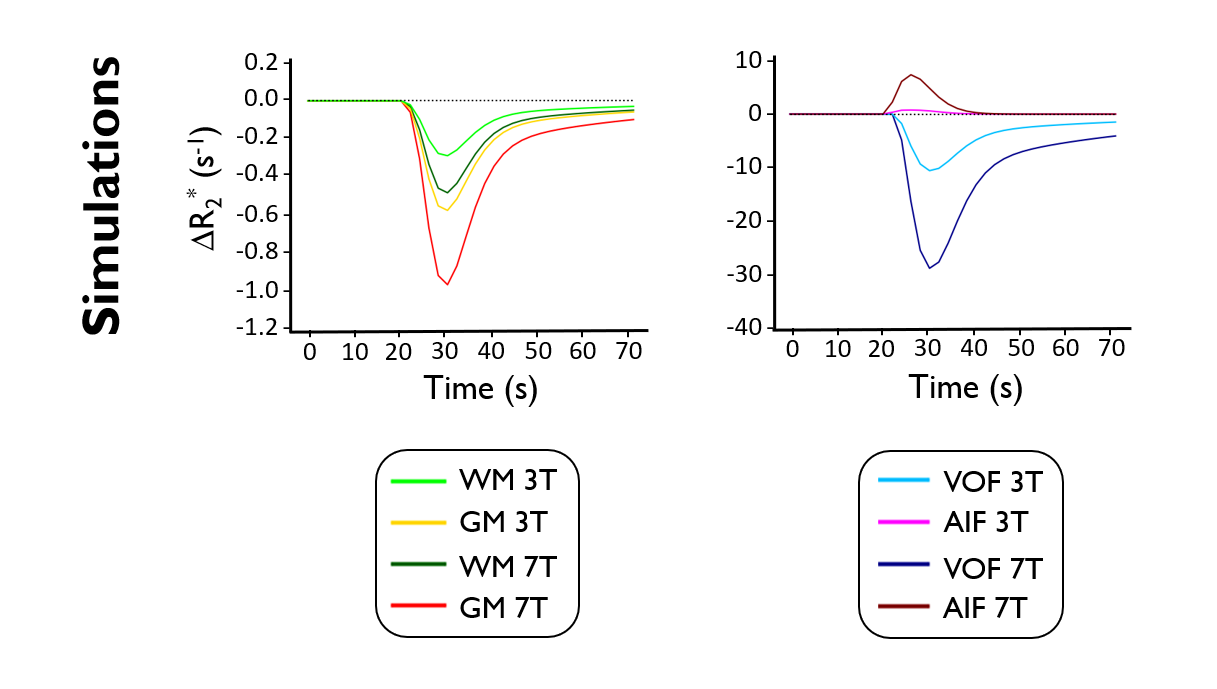

**Figure S2. Simulated Relaxation Rate Time Course Dynamics.** Simulated voxel configurations are based on parameters in Table S1.

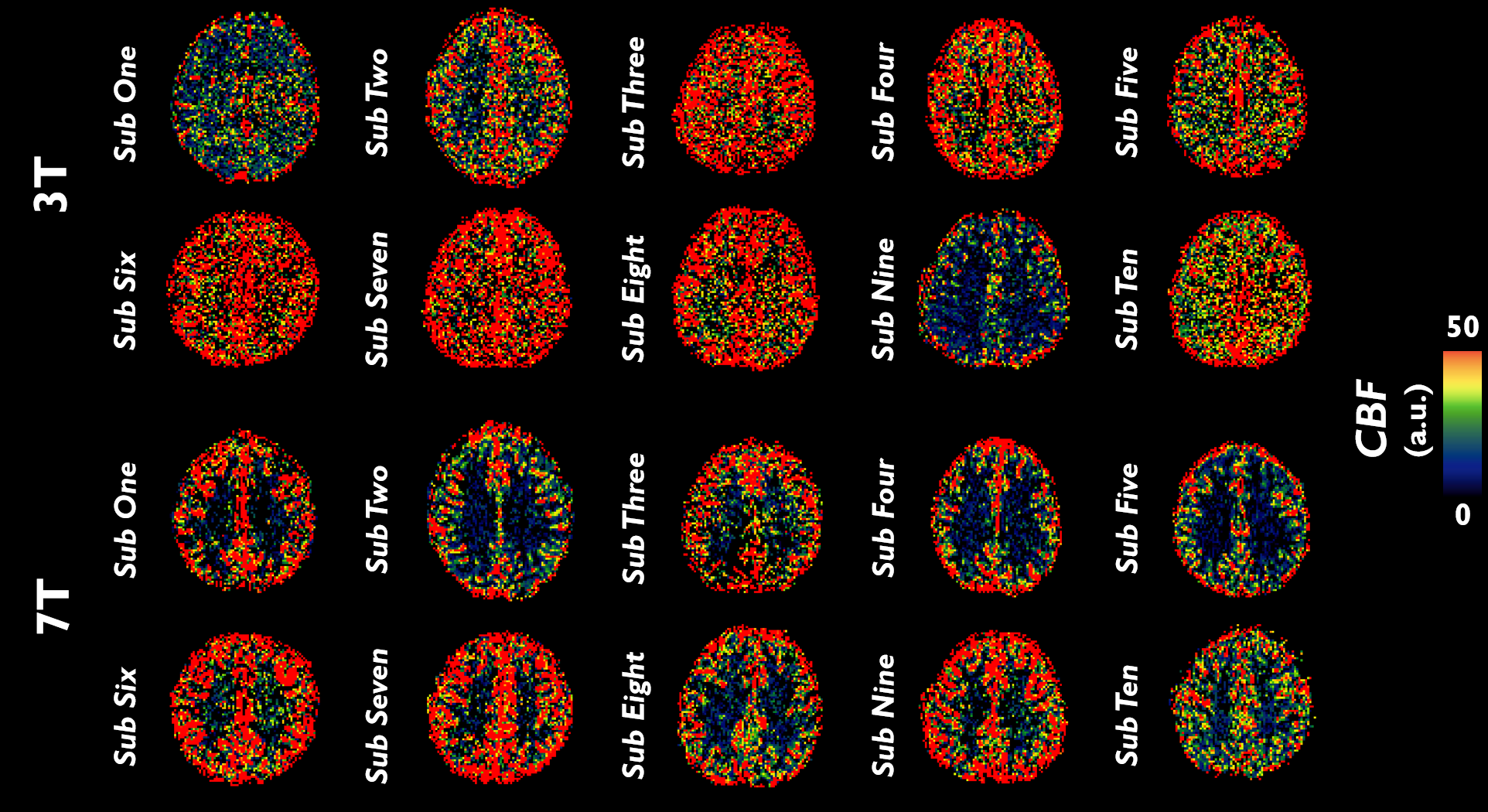

**Figure S3. Subject-wise *CBF* Perfusion Maps.** *CBF* (a.u.) maps are displayed for each subject at 3T and 7T. Axial views are just dorsal to the lateral ventricles.

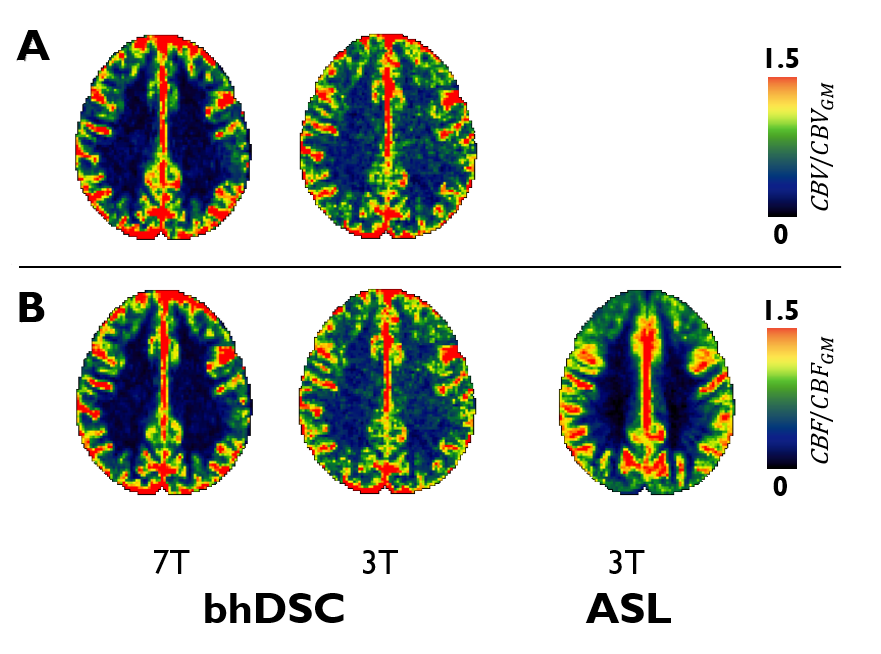

**Figure S4. Normalized *CBV* and *CBF*.** **A**. Subject-averaged *CBV* and **B.** *CBF* maps normalized to the average *CBV_GM_* and *CBF_GM_*, respectively, of an axial slice slightly dorsal to the lateral ventricles (z = 50) in MNI152 2 mm anatomical space. In comparison to GM, there are pronounced voxel-wise WM differences between 7T and 3T, but this is not unexpected given the low *CNR* in this anatomical region—our focus is on GM.

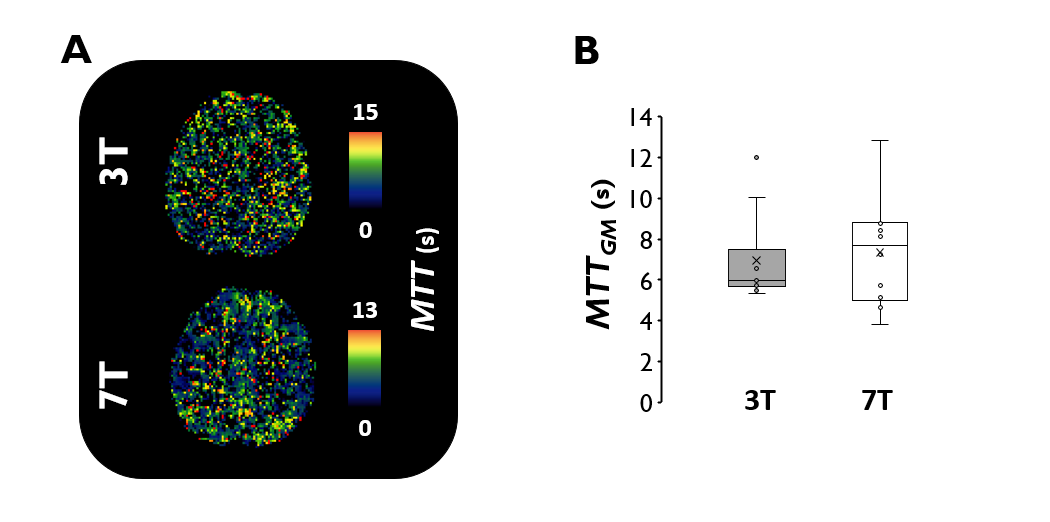

**Figure S5. bhDSC *MTT*. A.** *MTT* (s) map for a characteristic subject at 3T and 7T (axial view is slightly dorsal to lateral ventricles). **B.** Box and whisker plots illustrate the calculated *MTT_GM_* at 3T and 7T.

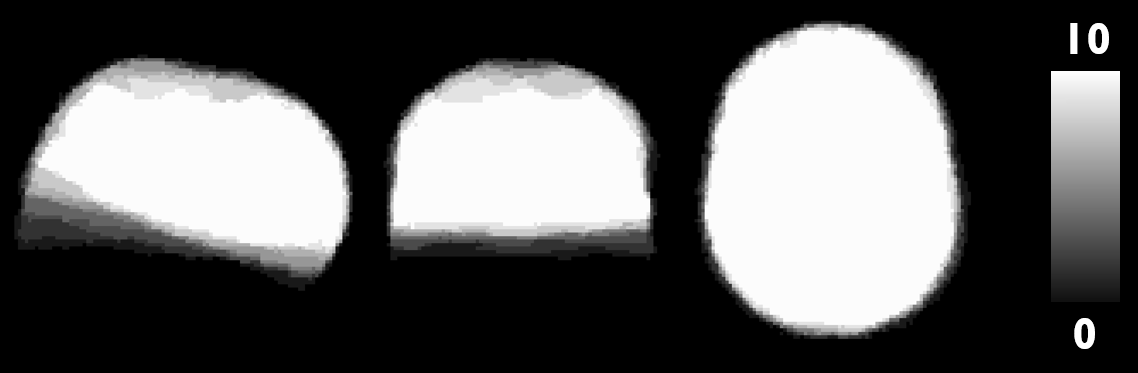

**Figure S6. ASL Standard Space Mask.** The ASL mask in MNI152 2 mm anatomical space was created based on where there was overlapping coverage amongst all (n = 10) participants (i.e., where mask value is equal to 10).

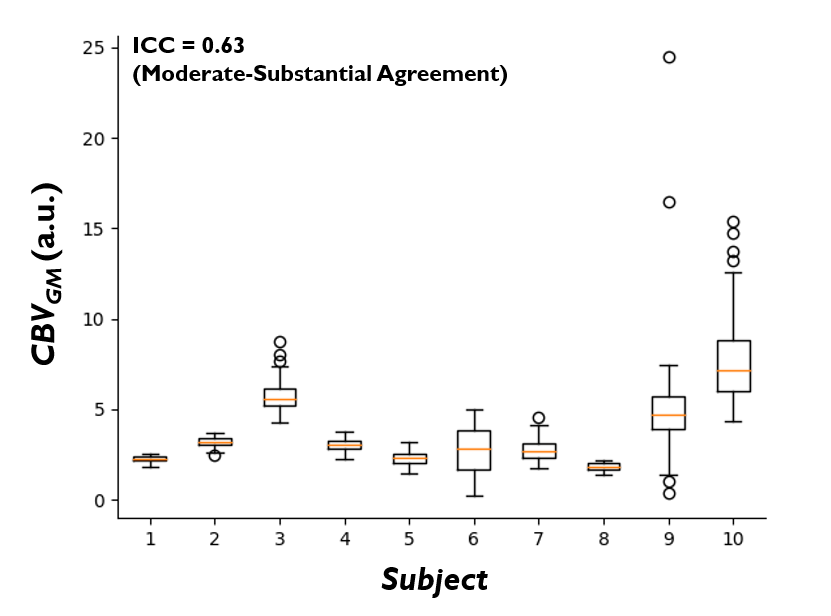

**Figure S7. Intra-Subject Precision of GM *CBV*.** Box and whisker plot illustrating GM *CBV* for 70 bootstrapped bolus combinations per subject at 7T (i.e., averaging of 4 randomly selected boluses for a total of 70 bolus combinations (_8_C_4_ = 70)). For intraclass correlation (ICC), refer to Methods.

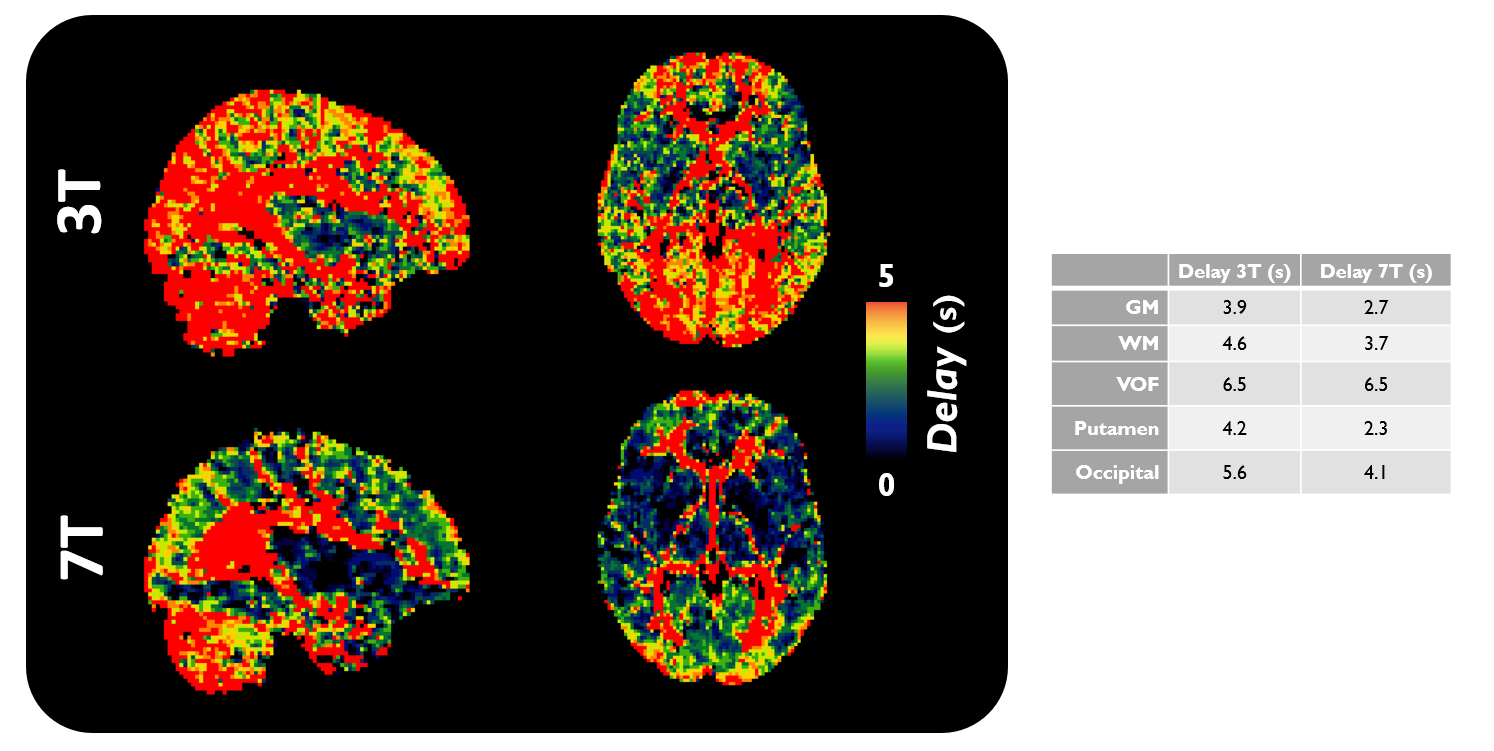

**Figure S8. bhDSC Delay Maps. Left.** Tissue delay maps relative to the AIF are illustrated in MNI152 2 mm anatomical space (sagittal and axial views are shown at 3T and 7T). **Right.** Calculated average delay values from various cerebral territories (note that masks for occipital GM and putamen are from the MNI structural atlas).^[89]^

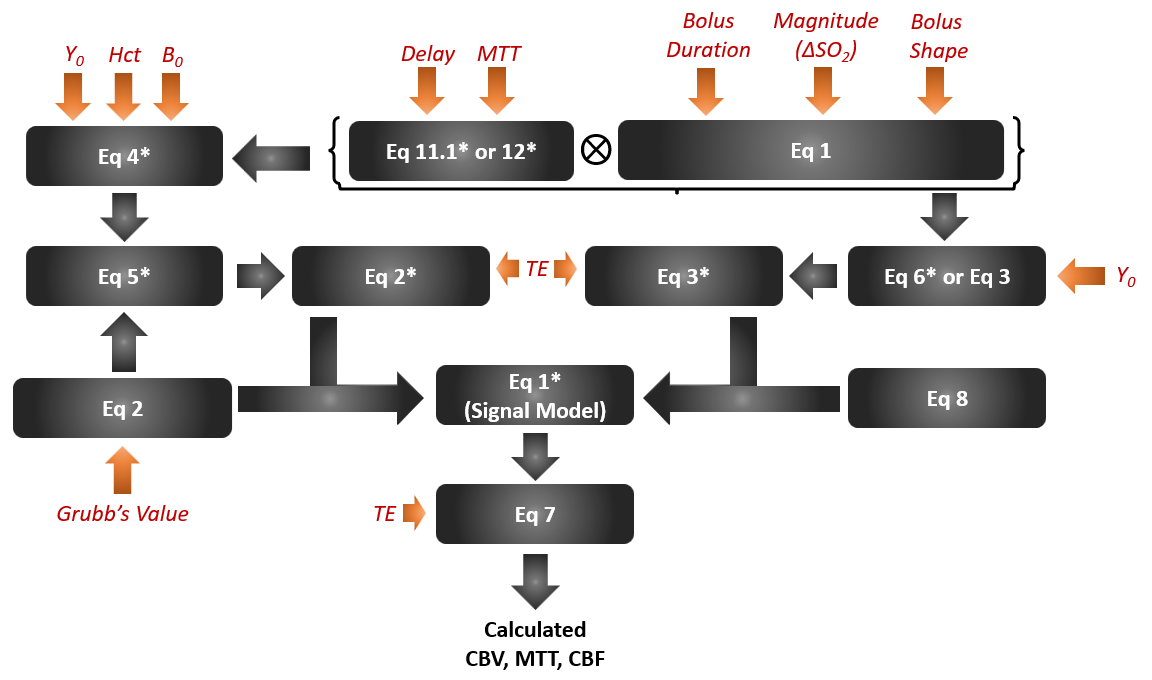

**Figure S9. Visual Illustration of Workflow for Signal Model.** Black arrows indicate where one equation’s output becomes the input for another equation. Parameters, shown in red, are based on values found in Table S1 and are described in the Simulation Methods section. This pipeline was used to simulate the signal and relaxation time courses for arterial, tissue, and venous voxels. The asterisk denotes when an equation is found in Schulman *et al.,* 2023.

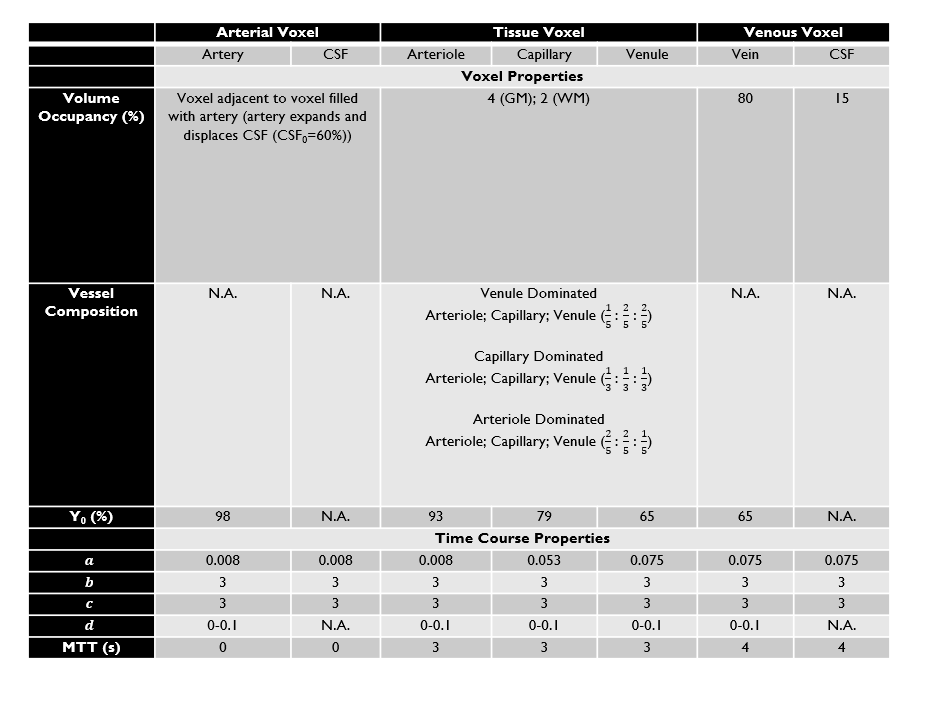

**Table S1. Summary of Simulated Time Course and Voxel Properties for Arterial, Tissue, and Venous Voxels.** The values in this table can be used for any arbitrary set of parameters, but these are the parameters used in our simulations. Note that any remaining volume occupancy, not filled by vessel or CSF, is composed of extravascular space.
